# A novel and robust method for assessing mitochondrial (dys)function in healthy and diseased frozen cardiac tissue

**DOI:** 10.1101/2024.12.11.627239

**Authors:** Liangyu Hu, Alexia van Rinsum, Rocco Caliandro, Xi Qi, Shuxiu Fan, Xinyi Jiang, Jur Massop, Melissa Bekkenkamp-Grovenstein, Christiane Ott, Tilman Grune, Melissa A.E. van de Wal, Werner J.H. Koopman, Marcel Giesbers, Monika Gladka, Jaap Keijer, Deli Zhang

## Abstract

Cardiovascular diseases are often associated with impairment in mitochondrial function detected by reduced mitochondrial oxygen consumption using high-resolution respirometry. However, existing respirometry protocols are limited by the necessity for fresh tissue samples. This study developed a method with tailored substrate-inhibitor titration (TSIT) of mitochondrial electron transport complexes (ETC) to measure mitochondrial function in frozen cardiac samples using high-resolution respirometry. Briefly, acetyl-CoA was added to fuel the tricarboxylic acid (TCA) cycle for NADH production, enabling complex I (CI)-linked respiratory assessment. NADH was then added to measure maximum CI-linked respiratory capacity, followed by rotenone and succinate to assess complex II (CII)-linked respiratory capacity. TSIT detected mitochondrial functional differences between frozen atrial and ventricular tissue, with comparable results as measured in fresh samples. It also detected cardiac mitochondrial dysfunction across various (patho)physiological mouse models (including aging, ischemia reperfusion, obesity, and CI deficiency) as well as in frozen human donor samples, highlighting its clinical potential. Furthermore, we showed the first evidence for supercomplexes (SCs) formation between ETC-SCs and the TCA cycle metabolon, underpinning TSIT feasibility. In conclusion, we established a novel, robust, sensitive and translational method (TSIT) for assessing mitochondrial (dys)function in frozen cardiac samples from various species, enabling flexible analysis of mitochondrial function in both laboratory and clinical settings.

## Introduction

Among all organs in the body, the heart has the highest metabolic rate reflecting its extensive workload and energy demand.^1^ Mitochondria generate the majority of energy needed for the activities of the heart^2^ and are thus crucial organelles for the beating heart. The energy is provided by mitochondrial respiration, which involves a series of redox reactions in the four mitochondrial inner membrane (MIM) electron transport complexes (ETC) followed by subsequent generation of adenosine triphosphate (ATP) by complex V (CV).^3^ The ETC consists of complex I to IV (CI-CIV) and several associated protein complexes, such as cytochrome c (CYTC), a loosely bound MIM protein that transfers electrons between CIII and CIV. The ETC complexes together with CV constitute the oxidative phosphorylation (OXPHOS) system. The continuous beating heart requires optimal mitochondrial OXPHOS function to sustain its high rate of ATP demand.

Impaired mitochondrial oxygen consumption or respiration, reflecting mitochondrial dysfunction, is involved in a wide range of cardiometabolic disorders^4–6^ and in cardiac aging.^7, 8^ Deficiency in mitochondrial CI in the *Ndufs4^−/−^* mice causes severe mitochondrial disorders and accelerated pressure overloaded heart failure (HF).^9, 10^ In agreement, a reduced enzymatic activity of CI^11^ and defects in CI-linked respiration were observed in human HF.^12–14^ Similarly, CI-mediated and fatty acid-mediated mitochondrial respiration as well as CI enzymatic activity were compromised in diabetic subsarcolemmal mitochondria^15^ and atrial myofibers from type 2 diabetic patients with diabetic cardiomyopathy.^16, 17^ This finding was corroborated by data from animal studies. For instance, a study using New Zealand obese (NZO) mice, characterized by severe obesity, insulin resistance, and a predisposition to type 2 diabetes, revealed a notable cardiac dysfunction and an impaired mitochondrial respiratory function in cardiac tissue under high-fat diet (HFD) feeding.^18^ Furthermore, the activity of mitochondrial ETC complexes, particularly of CI and CIV, declined with age in rat hearts.^7^ Mice subjected to coronary ischemia reperfusion (IR) were reported to develop cardiac mitochondrial dysfunction.^6, 19^ Collectively, the close association between cardiac diseases and compromised mitochondrial respiration underscores the importance of assessing mitochondrial respiratory function of cardiac tissue, which would reveal specific functional deficits and provide therapeutic guidance.

Currently, mitochondrial respiration of cardiac tissue is typically measured using mitochondrial preparations from freshly sampled tissue.^20^ The requirement for fresh tissue presents a severe limitation, since the analysis has to be performed shortly after sampling^21^, which requires a stringent alignment in time and proximity of physician and the specialized analysis lab. This introduces both study limitations and experimental variation due to compromised standardization. In addition, frozen cardiac tissue samples stored in patient biobanks cannot be used, limiting translational applications.^22^ Therefore, a robust method tailored for measuring mitochondrial respiration in frozen cardiac tissue to overcome these limitations is urgently needed.

Frozen samples have lost the coupling between the ETC and the ATP generation capacity due to the disruption of the MIM integrity during the freeze-thaw cycle. Therefore, many efforts have been dedicated to optimizing cryopreservation of biopsies to maintain the mitochondrial membrane integrity.^23–26^ However, cryopreservation is a delicate process, making it impractical for sample storage in clinical practice. Evidence from both the spectrophotometric assays of individual ETC complex activities^27^ as well as the functional analysis of mitochondrial supercomplexs (SCs, e.g. the I-II-III_2_-IV_2_ SC)^28^ indicated that the electron transport activity is maintained in frozen tissue samples despite the uncoupled condition. In respirometry, the uncoupled respiratory state is assessed by the addition of a chemical uncoupler to gain insight in the maximal ETC capacity. A reduction in the maximal respiratory capacity may limit the ability to cope with stressors, resulting in mitochondrial dysfunction.^29^ The situation in frozen samples represents the condition in which uncoupled maximal respiration is assessed. By directly supplying ETC substrates as well as loosely attached ETC components, such as CYTC, the respiratory capacity that is comparable to uncoupled maximal respiration in the conventional respirometry using fresh tissue would be possible in frozen tissue. Indeed, two methods were recently explored to measure mitochondrial CI respiratory capacity in frozen patient muscle biopsies using the Oroboros O2k^30^ and in various tissues using the Seahorse XF Analyzer.^22^ However, a robust method using high-resolution respirometry to assess cardiac ETC driven mitochondrial respiration in frozen tissue, capable of detecting respiratory activities of multiple mitochondrial complexes within a single round of measurement, is not yet available.

Compared to the Seahorse XF Analyzer, the Oroboros O2k allows for detailed measurement of mitochondrial respiration mediated by various mitochondrial complexes within a single round. Therefore, we aimed to develop a method, using the Oroboros O2k, to measure mitochondrial respiratory capacities in frozen cardiac tissues from both mice and human by Tailored Substrate-Inhibitor Titration of mitochondrial complexes (named TSIT). Frozen cardiac tissues from various physiological and pathophysiological mouse models and human donors with known mitochondrial dysfunction were used to further test and validate the potential and robustness of the application of TSIT.

## Methods

All animal procedures required were ethically approved by the Animal Welfare Committee of Wageningen University regarding the mice used for atrial and ventricular tissue comparison (2020.W-0019.008) and the ageing mice (2020.W-0019.007), by the Radboud University Nijmegen regarding the *Ndufs4*^−/−^ mice (2017.W-0017-007), by the German Institute of Human Nutrition regarding the NZO mice (2347-46-1029), or by Amsterdam Medical Center regarding the IR mice (AVD11800202114603). Human cardiac samples were collected and characterized at the Amsterdam Medical Center (Table S1). The study conformed to the principles of the Declaration of Helsinki. The institutional ethical review board approved the study (2024.0643), and patients gave written informed consent.

### Cardiac tissue collection from mice and human patients

All mice had free access to food and water and were maintained at the standard temperature and humidity of the respective animal facilities. To compare the difference in mitochondrial respiration between atrial and ventricular tissues, cardiac tissue samples were collected from 6-month-old BL6JRcc(BL6J)-*Nnt^+^*/Wuhap (*Nnt*^wt^) mice fed with standard chow, an in-house bred mouse line similar to C57BL/6JRccHsd. The mice were killed by decapitation and dissected immediately. After washing in ice-cold phosphate-buffered saline (PBS), the heart was split into right atrium (RA), left atrium (LA), right ventricle (RV) and left ventricle (LV). The samples were then either immediately snap frozen in liquid nitrogen and stored at −80 °C before the measurement (n=6), or freshly immersed in the BIOPS buffer (2.77 mM CaK_2_ EGTA,7.23 mM K_2_EGTA, 5.77 mM Na_2_ATP, 6.56 mM MgCl_2_, 20 mM taurine, 15 mM Na_2_ phosphocreatine, 20 mM imidazole, 0.5 mM DTT, 50 mM MES, pH 7.1) and proceeded for respiration measurement within 6 h (n=5).

Frozen cardiac tissue samples from one physiological mouse model and three pathophysiological mouse models were utilized to evaluate the sensitivity and applicability of TSIT. In detail, the physiological model of natural ageing compared young (4-month-old) and old (23-month-old) mice (*Nnt*^wt^, n=6 each), fed with standard chow diet. The pathophysiological models included: 1) chow fed C57BL/6N mice subjected to myocardial infarction surgery to mimic IR and sham controls (n=6 each);^31^ and 2) C57BL/6J mice fed with a standard chow diet (BL/6J control) and the NZO mice fed with a standard chow diet (NZO+SD) and a 35 en% high-fat diet (NZO+HFD), respectively (n=6 each);^18^ and 3) chow fed *Ndufs4*^−/−^ mice and the control wildtype (WT) mice (n=6 each).^32^ LV samples from mouse models above were snap frozen and stored in −80 °C until further analysis.

Frozen human LV free-wall tissue samples from human donors with cardiac pathology (n=5) or with no/low level cardiac pathology (n=4) were measured to prove the utility of TSIT in clinical practice. The detailed information about the human donors is listed in Table S1.

### Preparation of fresh cardiac tissues with mechanical permeabilization for high-resolution respirometry

Fresh mouse atrial and ventricular tissue samples were kept in ice-cold BIOPS buffer for a maximum of 6 h. For subsequent respirometry analyses, the tissue was quickly dried on filter paper and weighed before being transferred to a 1.5 ml Eppendorf tube with 50 μl of ice-cold BIOPS buffer. The average tissue mass varied between 0.4-0.7 mg. Mechanical permeabilization was performed using a hand-held tissue homogenizer (47747-370, VWR) with a disposable pestle (431-0094, VWR) at the speed of 12000 RPM. To ensure intact mitochondrial membranes in the fresh cardiac tissue, 3 s of mechanical permeabilization for atrial tissue and 1 s for ventricular tissue were used. The procedures were performed on ice throughout the experiment. Fresh tissue preparations were then loaded into the chambers of an Oroboros O2k (series H-0391, Oroboros Instruments) with 2 ml mitochondrial respiration medium MiR05 (6010101, Oroboros Instruments) containing 0.5 mM EGTA, 3 mM MgCl_2_, 60 mM lactobionic acid, 20 mM taurine, 10 mM KH_2_PO_4_, 20 mM HEPES, 110 mM D-sucrose, and 1 g/L BSA (pH 7.1).

### Preparation of frozen cardiac tissue homogenates for high-resolution respirometry

Frozen cardiac tissue was quickly thawed in ice-cold BIOPS buffer, dried on filter paper and weighed before being transferred to a 1.5 ml Eppendorf tube with 50 μl of ice-cold BIOPS buffer. The average tissue mass varied between 0.4-0.7 mg for frozen mouse atrial and ventricular tissue, and 1.8-2.2 mg for frozen human ventricular tissue. Tissue was homogenized using a hand-held tissue homogenizer (47747-370, VWR) with a disposable pestle (431-0094, VWR) at the speed of 12000 RPM until no visible tissue debris was observed. The optimal homogenization time of frozen tissue was approximately 5 s for mouse ventricular tissue, 10 s for mouse atrial tissue, and 15 s for human ventricular tissue. The procedures were performed on ice throughout the experiment. 50 μl frozen tissue homogenates with or without centrifugation at 1000 x *g* for 5 min at 4 °C were then loaded into the chambers of an Oroboros O2k (series H-0391, Oroboros Instruments) with 2 ml mitochondrial respiration medium MiR05 (6010101, Oroboros Instruments).

### High-resolution respirometry of fresh cardiac tissue

High-resolution respirometry (Oroboros O2k, series H-0391) using mechanically permeabilized fresh cardiac tissue was performed at 37 °C in a hyper oxygenated condition (300-450 μM O_2_), allowing for a sufficient oxygen concentration gradient into the mechanically permeabilized fresh tissue. Mitochondrial respiration was assessed by the conventional substrate-uncoupler-inhibitor titration (SUIT) protocol with minor modifications as follows (SUIT-002, https://wiki.oroboros.at/index.php/SUIT-002, *Figure 1A*): All concentrations given, except carbonyl cyanide-p-trifluoromethoxyphenylhydrazone (CCCP), are the final concentrations. Briefly, 7.5 mM ADP was added to stimulate the consumption of endogenous fuel-substrates without ADP limitation (OXPHOS state). 0.3 mM malate and 0.5 mM octanoylcarnitine were added to measure OXPHOS respiration through the fatty acid oxidation (FAO) pathway. 10 µM Cytc was then added to test mitochondrial outer membrane integrity. Samples that responded to Cytc addition by a 20% increase in respiration were excluded. Then, 2 mM malate, 5 mM pyruvate and 10 mM glutamate were added to measure OXPHOS respiration through FAO and CI (FAO+CI), followed by addition of 10 mM succinate to measure OXPHOS respiration through FAO, CI, and CII (FAO+CI+CII). Next, titrations with 1 μl of 0.1 μM CCCP were used to measure the uncoupled maximal electron transport chain capacity (ETS state). In the end, 1 µM rotenone and 5 μM antimycin A were added to measure residual oxygen consumption (Rox) or non-mitochondrial respiration. Mitochondrial respiration calculated as oxygen (O_2_) flux subtracted from Rox, was normalized to wet tissue weight and expressed as pmol/s*mg tissue.

**Fig. 1:**
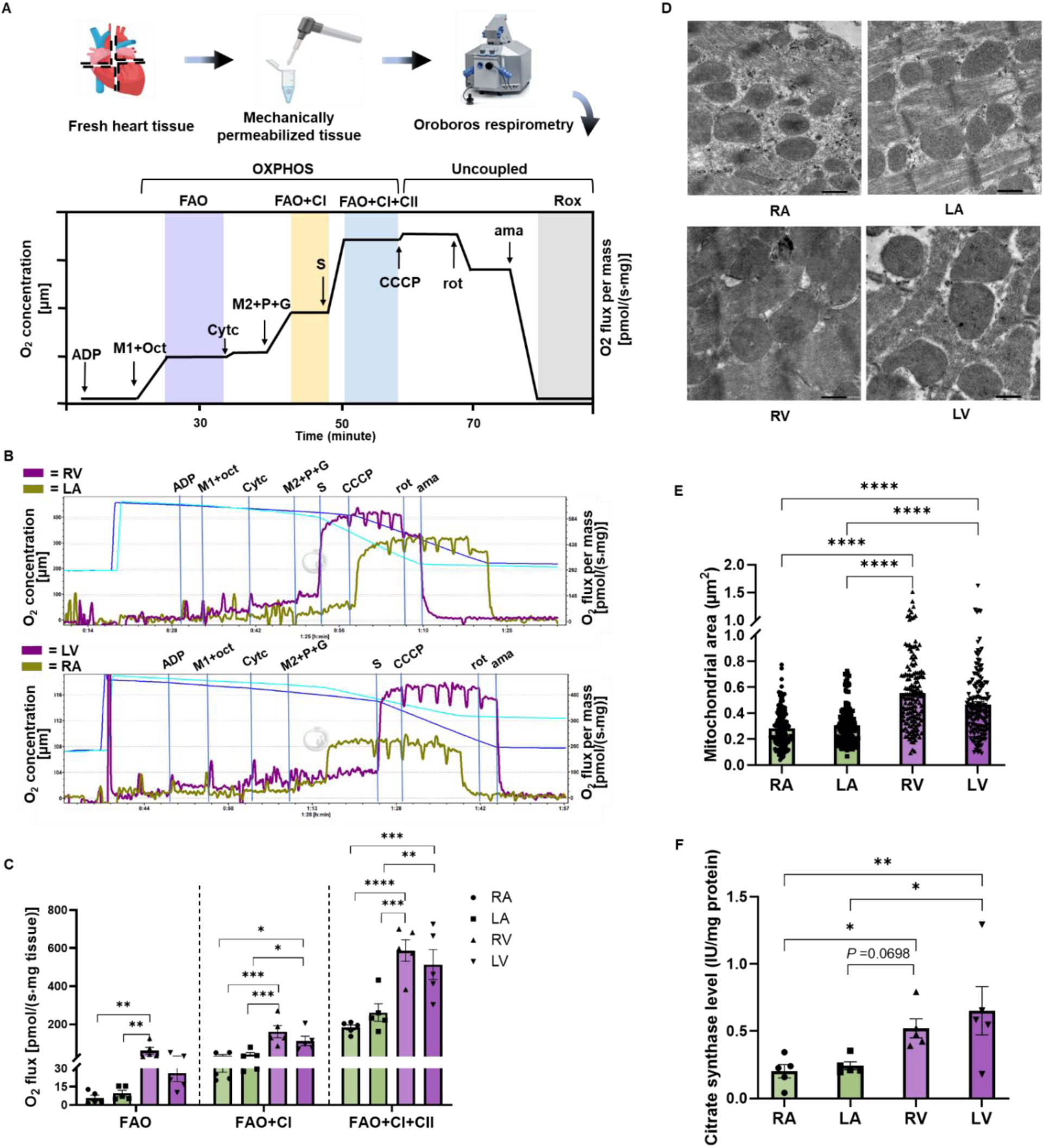
Mitochondrial respiration and ultrastructure in freshly harvested mouse cardiac tissue. (*A*) Schematic representation of the sample preparation and respirometry protocol. Fresh dissected mouse cardiac tissues from right atrium (RA), left atrium (LA), right ventricle (RV), left ventricle (LV) were weighed, homogenized and immediately measured using high-resolution respirometry according to the SUIT-002 protocol as illustrated. Fatty acid oxidation-boosted respiration (FAO, purple) was determined at 7.5 mM ADP, 0.3 mM malate (M1) and 0.5 mM octanoylcarnitine (oct). 10 µM cytochrome c (Cytc) was then added to test mitochondrial outer membrane integrity. FAO+complex I (CI)-linked respiration (FAO+CI, yellow) was determined at an additional 2 mM malate (M2), 5 mM pyruvate (P) and 10 mM glutamate (G). FAO+CI+complex II (CII)-linked respiration (FAO+CI+CII, blue) was determined at an additional 10 mM succinate (S). FAO+CI+CII-linked uncoupled maximal respiration was determined at an additional titration of 1 μl of 0.1 μM carbonyl cyanide-p-trifluoromethoxyphenylhydrazone (CCCP). Residual oxygen consumption (Rox, gray) was determined at an additional 2µM rotenone (rot) and 5 μM antimycin A (ama). (*B*) Representative traces of fresh RA, LA, RV, and LV tissues from BL6JRcc(BL6J)-*Nnt^+^*/Wuhap mice. Purple traces indicate the ventricular O_2_ flux and olive-green traces indicate the atrial O_2_ flux. (*C*) Quantification of FAO-, FAO+CI- and FAO+CI+CII-linked mitochondrial respiratory capacities in fresh atrial and ventricular tissues (n=5). (*D*) Representative images of mitochondrial morphology in RA, LA, RV and LV tissues, scale bar=500 nm. (*E*) Quantification of mitochondrial size in atrial and ventricular tissues (n=150 mitochondria from three biological replicates). (*F*) Citrate synthase levels, an indicator of mitochondrial mass, in fresh RA, LA, RV, and LV tissues (n=5). Data are expressed as mean ± standard error of the mean. One-way analysis of variance (ANOVA) followed with a post-hoc Fisher’s LSD test was used. \**P* < 0.05, \*\**P* < 0.01, \*\*\**P* < 0.001, \*\*\*\**P* < 0.0001.

### High-resolution respirometry of frozen cardiac tissue

High-resolution respirometry of frozen cardiac tissue homogenate using TSIT (Oroboros O2k, series H-0391) was performed in a normal oxygenated condition (80-180 μM O_2_) since oxygen easily permeated into the frozen tissue homogenate. The sequential titration of the substates and inhibitors of mitochondrial complexes according to TSIT, given in final concentrations (unless otherwise stated), was as follows (*Figure 2A*): 2 mM malate, 100 µM NAD^+^ and 10 µM Cytc were added to determine ‘Basal’ respiratory capacity. Next, 40 µM palmitoyl-CoA, 40 µM FAD, and 1.2 mM coenzyme A (CoASH) were added to determine FAO-linked respiratory capacity. 150 µM acetyl-CoA (A2181, sigma) served to boost the tricarboxylic acid (TCA) cycle to produce NADH from NAD^+^ for the assessment of the CI-linked respiratory capacity. Then, to surpass the possible limitation of the CI substrate NADH supplied by the TCA cycle, 20 μl of 10 mM NADH was titrated to determine the maximum CI-linked respiration until the flux was stable. Subsequent addition of 1 µM rotenone and titration of 20 µl of 1 M succinate were used to assess CII-linked respiratory capacity. Finally, Rox was determined after the sequential addition of 5 mM malonate (inhibitor of CII) and 2.5 µM antimycin A. For each biological sample at least 2 technical replicates were performed. Mitochondrial respiration calculated as O_2_ flux subtracted from Rox, was normalized to tissue wet weight and expressed as pmol/s*mg tissue.

**Fig. 2:**
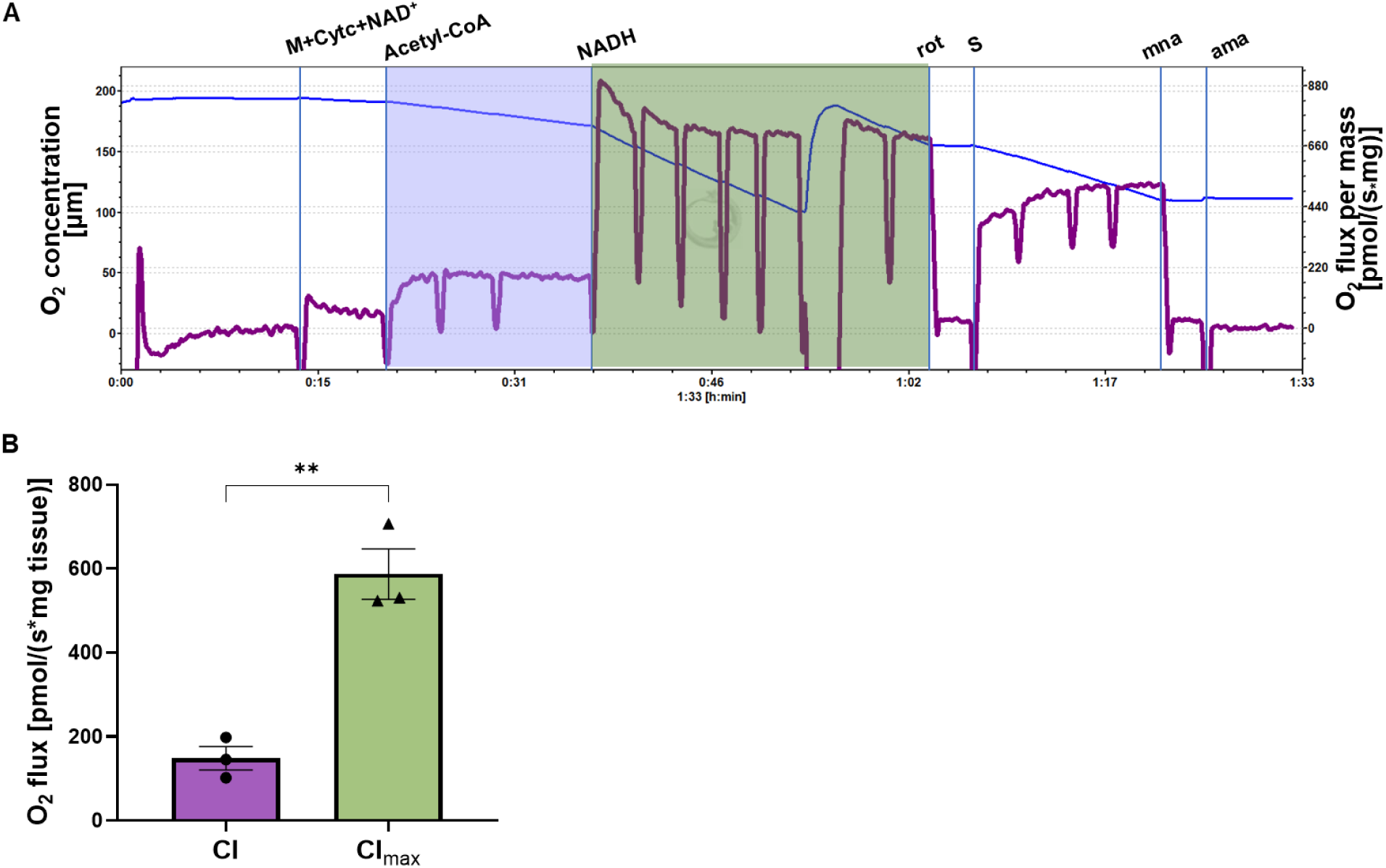
High-resolution respirometry measurement of complex I (CI)-linked respiratory capacity at physiological condition and at maximum capacity in frozen mouse cardiac tissue using the TSIT method. (A) Representative trace of frozen left ventricular (LV) tissue. Basal respiratory capacity was determined at 2 mM malate (M), 100 µM NAD^+^ and 10 µM cytochrome c (Cytc). CI-linked respiratory capacity (purple) was determined at titration of 15 µl of 20 mM acetyl-CoA to mimic the physiological condition. Maximum CI-linked respiratory capacity (CI_max_, olive-green) was determined with titration of 20 μl of 10 mM NADH. Complex II-linked respiratory capacity was determined at 1 µM rotenone (rot) and titration of 20 µl of 1 M succinate (S). Mitochondrial respiration was calculated as O_2_ flux subtracted from residual oxygen consumption, which was determined at an additional 5 mM malonate (mna) and 5 µM antimycin A (ama). (B) Quantified O_2_ flux showed that the CI_max_ (olive-green) was roughly 4 times that of the CI-linked respiratory capacity (purple) mediated by acetyl-CoA in frozen LV tissue (n=3). Data are expressed as mean ± standard error of the mean. A two tailed unpaired Student t-test was used. \*\**P* < 0.01. \*\**P* < 0.01.

### Citrate synthase activity

Citrate synthase (CS) activity, an indicator of mitochondrial mass,^33^ was determined using the CS assay kit according to the manufacturer’s instructions (CS0720, Sigma). Briefly, the corresponding frozen tissue from the same animal used for high-resolution respirometry measurement was weighed and homogenized in lysis buffer (150 μl lysis buffer per mg tissue) with the hand-held tissue homogenizer (47747-370, VWR). The lysis buffer contained 50 mM Tris-HCl pH 7.4, 150 mM NaCl, 1% Triton X-100, 1 mM ethylenediaminetetraacetic acid, and protease (04693159001, Roche) and phosphatase inhibitor cocktail (04906837001, Roche). After a freeze-thaw cycle (−80 °C freezing for 10 min, 4 °C thawing), the homogenate was centrifuged at 10,000 x *g* for 5 min at 4 °C to remove cell debris, and 4 μl of supernatant was used to measure CS activity based on the real-time absorbance kinetics at 412 nm in the linear range with an interval of 15 s for 3 min using the plate reader (235197, BioSPX). The CS activity (IU/ml) was corrected by the tissue protein concentration (mg protein/ml) and expressed as IU/mg protein.

### Transmission electron microscopy

Atrial and ventricular tissue samples were cut into pieces of about 1mm^3^ and fixed in a mixture of 2% glutaraldehyde (16316-10, EMS) and 2% formaldehyde (15700, EMS) in 0.1 M phosphate/citrate buffer overnight at 4 °C. Tissue was then washed with 0.1 M phosphate/ citrate buffer and subsequently postfixed in 1% osmium tetroxide (19130, EMS) for 1 h at room temperature, washed with deionized water, dehydrated in ethanol (30%, 50%, 70%, 80%, 90% and 100% for 5-10 minutes each), embedded in Spurr resin (14300, EMS) overnight, and polymerized at 70°C for 8 h. Ultrathin sections (60 nm) were stained with 2% uranyl acetate (22400, EMS) and 3% lead citrate (22410, EMS) for 10 minutes, and then viewed on copper formvar coated grids under a transmission electron microscope (JEM 1400 Plus Transmission EM). From images of three biological replicates, 150 mitochondria of each group were analyzed. Mitochondrial size and number were analyzed using Fiji software (ImageJ 1.52j).

### Separation of mitochondrial supercomplexes by blue-native gel electrophoresis

Frozen mouse cardiac tissue (around 10-15 mg) was homogenized with a hand-held homogenizer (47747-370, VWR) with a disposable pestle (431-0094, VWR) in 1 ml ice-cold mitochondrial lysis buffer consisting of 250 mM sucrose, 20 mM Tris-HCl pH 7.4, 1 mM ethylenediaminetetraacetic acid, and a protease inhibitor cocktail (04693159001, Roche). The tissue homogenate was freed from cells debris and nuclei by centrifugation at 1000 x *g* for 10 min at 4 °C. The supernatant was then centrifuged at 10,000 x *g* for 10 min at 4 °C to obtain a mitochondrial enriched pellet, which was resuspended in 100 µl fresh mitochondrial lysis buffer. The protein concentration was measured using DC protein assay kit according to the manufacturer’s instructions (500-0116, Bio-Rad). Then, 60 µg of mitochondria-enriched fraction protein was pelleted at 20,000 x *g* for 10 min at 4 °C and then resuspended in 12 µl solubilization buffer (50 mM NaCl, 50 mM imidazole-HCl, 5 mM 6-aminocaproic acid, 1 mM EDTA, pH 7.0). Thereafter, 540 µg digitonin was added, and incubated on ice for 10 min for further solubilization. Then, the solubilized samples were centrifuged at 20,000 x *g* for 20 min at 4 °C and supernatant was mixed with 60 µg Coomassie Brilliant Blue G before loading (27815, sigma). NativeMark unstained protein standard was included (LC0725, Invitrogen). Blue native PAGE analysis was performed using a NativePAGE 3-12% Bis-Tris gel (BN2011BX10, Invitrogen) and electrophoresis was done at 50 V for 30 min and then 150 V for 120 min in XCell SureLock system (EI0001, Novex). Protein was transferred to a methanol-activated PVDF membrane (IPFL85R, sigma) at 30 V overnight at 4°C (1703930, Bio-Rad). The membrane was then washed with 1x TBST with 2% SDS, pH 7.6 at 60 °C, blocked with Intercept Blocking Buffer (927-60001, LI-COR) for 1.5 h at room temperature, and incubated with primary antibodies anti-CI (A21344, Invitrogen, 1:1000 dilution), anti-succinate dehydrogenase A (SDHA, ab14715, Abcam, 1:1000 dilution), anti-CIV (ab16056, Abcam, 1:1000 dilution), or anti-CS (14309S, Cell signaling, 1:1000 dilution) overnight at 4°C. The membrane was then incubated with secondary antibodies IRDye 800CW Donkey anti-mouse (926-32212, LI-COR, 1:5000 dilution) or IRDye 680RD Donkey anti-rabbit (926-68073, LI-COR, 1:5000 dilution) for 1 h at room temperature and visualized using an Odyssey scanner (LI-COR).

### Protein extraction and Western blotting analysis

Frozen patient left ventricular tissue was lysed in radioimmunoprecipitation assay (RIPA) buffer containing 10 mM Tris-base pH 7.6, 150 mM NaCl, 0. 5% sodium deoxycholate, 1% SDS, 1% Triton X-100, and protease (04693159001, Roche) and phosphatase inhibitor cocktail (04906837001, Roche). 8 μg protein per lane was separated on a 4-12% NuPage gel (NP0323BOX, Invitrogen) and electrophoresed at 110 V for 30 min and then 150 V for 1 h in XCell SureLock system (EI0001, Novex). Proteins were transferred to a methanol-activated PVDF membrane (IPFL85R, sigma) at 300 mA for 90 min at room temperature (1703930, Bio-Rad). The membrane was washed with 1x TBS for 5 min, blocked with Intercept Blocking Buffer (927-60001, LI-COR) for 1 h at room temperature, and incubated with primary antibodies anti-OXPHOS (ab110413, abcam, 1:1000 dilution) or anti-VDAC1 (ab235143, abcam, 1:1000 dilution) overnight at 4°C. The membrane was subsequently incubated with secondary antibodies IRDye 800CW Donkey anti-mouse (926-32212, LI-COR, 1:5000 dilution) or IRDye 680RD Donkey anti-rabbit (926-68073, LI-COR, 1:5000 dilution), respectively, for 1 h at room temperature and visualized using an Odyssey scanner (LI-COR).

### Statistics

Oxygen concentration and oxygen flux were recorded and processed by DatLab software 7.4 (Oroboros Instruments). To compare mitochondrial respiration, mitochondrial size and mass among ventricular and atrial tissues, as well as mitochondrial respiration in the NZO mouse model fed with and without HFD, one-way analysis of variance (ANOVA) followed with a post-hoc Fisher’s LSD test was used. To compare each mitochondrial complex-driven respiratory capacities between physiological or pathophysiological mouse and human samples and their respective controls, a two tailed unpaired Student t-test was used. To correlate the mitochondrial respiratory capacity and relative complex protein level, Pearson correlation analysis was used. The statistical significance was shown as \**P* < 0.05, \*\**P* < 0.01, \*\*\**P* < 0.001, \*\*\*\**P* < 0.0001. All the comparisons and graphs were made using the GraphPad Prism software (v9.5.1).

## Results

### High-resolution respirometry of fresh cardiac tissue reveals significant differences in mitochondrial respiration between atria and ventricles in mice

Mitochondrial respiration of fresh mouse cardiac tissues from both atria and ventricles was measured using high-resolution respirometry according to the SUIT-002 protocol (Oroboros O2k) (*Figure 1A*). We found that the FAO+CI-linked, as well as FAO+CI+CII-linked mitochondrial OXPHOS respiration were significantly higher for ventricles (RV and LV) than for atria (RA and LA) (*Figure 1B* and *C*). No significant difference between the atria (RA vs LA) or between the ventricles (RV vs LV) was found. No difference was observed between mitochondrial respiration in the OXPHOS state and the uncoupled maximal ETS state (*Figure S1*), which was in line with previous study that the ETS state was found to be similar to the value of ADP-stimulated respiration (OXPHOS states) in mouse tissue.^20^ Furthermore, we observed a larger mitochondrial size (*Figure 1D* and *E*) and mass (*Figure 1F*) in ventricular tissue than those in atrial tissue, which might explain the differences in mitochondrial respiration between atria and ventricles. Taken together, these data indicate evident differences in mitochondrial respiration and morphology between atrial and ventricular tissues in mice.

### Establishment and optimization of TSIT using mouse frozen cardiac tissue

Previously either frozen tissue homogenate without centrifugation^30^ or with low speed centrifugation^22^ were used in mitochondrial respirometry. To test the centrifugation effects in TSIT, we compared the mitochondrial respiration of frozen LV tissue homogenates with and without centrifugation at 1000 x *g* for 10 min using the same amount of tissue homogenate. We found that centrifugation significantly reduced the mitochondrial respiratory capacities (*Figure S2A* and *B*), indicating loss of mitochondria due to centrifugation prior to the measurement. In addition, after correcting the O_2_ flux to the protein concentration, a larger standard deviation in mitochondrial respiratory capacities was found in centrifuged samples compared with the uncentrifuged samples, although the significant differences between centrifuged and uncentrifuged samples were less abundant (*Figure S2C*). Therefore, whole tissue homogenate without centrifugation was applied in TSIT.

The conventional SUIT protocol using fresh tissue showed a smaller mitochondrial respiration through the FAO pathway compared to CI- and CII-linked respiration (*Figure 1C*). We explored whether FAO-linked respiratory capacity could be assessed in frozen LV tissue. However, this was not feasible when using theoretically relevant FAO substrates and components, specifically 40 µM palmitoyl-CoA, 40 µM FAD^+^, and 1.2 mM CoASH (*Figure S3*). In contrast, a substantial O_2_ flux after the subsequent addition of acetyl-CoA (CI-linked respiratory capacity) and succinate (CII-linked respiratory capacity) indicated that the acetyl-CoA-mediated TCA cycle and the mitochondrial electron transport capacity from respectively CI and CII to (CIII and) CIV were preserved in the frozen tissue (*Figure S3*). Therefore, we further focused on the CI- and CII-linked respiratory capacities.

Since the acetyl-CoA-mediated TCA cycle generated a limited amount of the CI substrate NADH, we then directly titrated NADH to measure the maximum potential of CI-linked respiratory capacity in frozen tissue. The results showed that the addition of NADH resulted in a flux that was roughly 4 times that of the CI-linked respiratory capacity meditated by acetyl-CoA in frozen moue LV tissue (*Figure 2*). Nevertheless, CI-linked respiration with addition of acetyl-CoA and the necessary substrates for TCA cycle better mirrors physiological conditions wherein the TCA cycle and the ETC are coupled for NADH production, and hence, we decided to measure the acetyl-CoA mediated CI-linked respiratory capacity in the following tests of mitochondrial function in various frozen cardiac samples using TSIT.

Furthermore, it is well established that the ETC complexes can form SCs^34–36^. Previous research suggested that TCA cycle enzymes can also form SCs, named metabolon^37, 38^. Moreover, a recent proteomic study showed that knockdown of CS, a key enzyme of the TCA cycle, significantly reduced the expression of mitochondrial ETC enzymes, especially the CI subunits.^39^ Since TSIT demonstrated that mitochondrial respiratory capacity can be assessed in frozen cardiac tissue with acetyl-CoA as substrate for TCA cycle, but not with substrates for FAO (*Figure S3*), we hypothesized that the TCA cycle enzymes might couple with the SCs of ETC in the frozen cardiac tissue. To test this hypothesis, we assessed mitochondrial SCs formation in frozen tissue homogenates from mouse LV tissues with blue native gel electrophoresis (BNGE). As expected, BNGE showed that CI and CIV formed SCs at a high molecular weight over 1M kDa with individual CI and CIV also being visible (*Figure 3A* and *S4*). Due to technical limitations of BNGE,^28^ CII and CIV individually were observed, while no CII-CIV SCs could be visualized as expected (*Figure 3B* and *S4*). Intriguingly, we found the interactions of CS with CI (*Figure 3C* and *S4*) and of CS with CII (*Figure 3D* and *S4*), indicating the TCA cycle enzymes indeed coupled with ETC complexes, which could explain why respiration measurement in frozen tissue homogenate using substrates of the TCA cycle (i.e. acetyl-CoA) could be achieved.

**Fig. 3:**
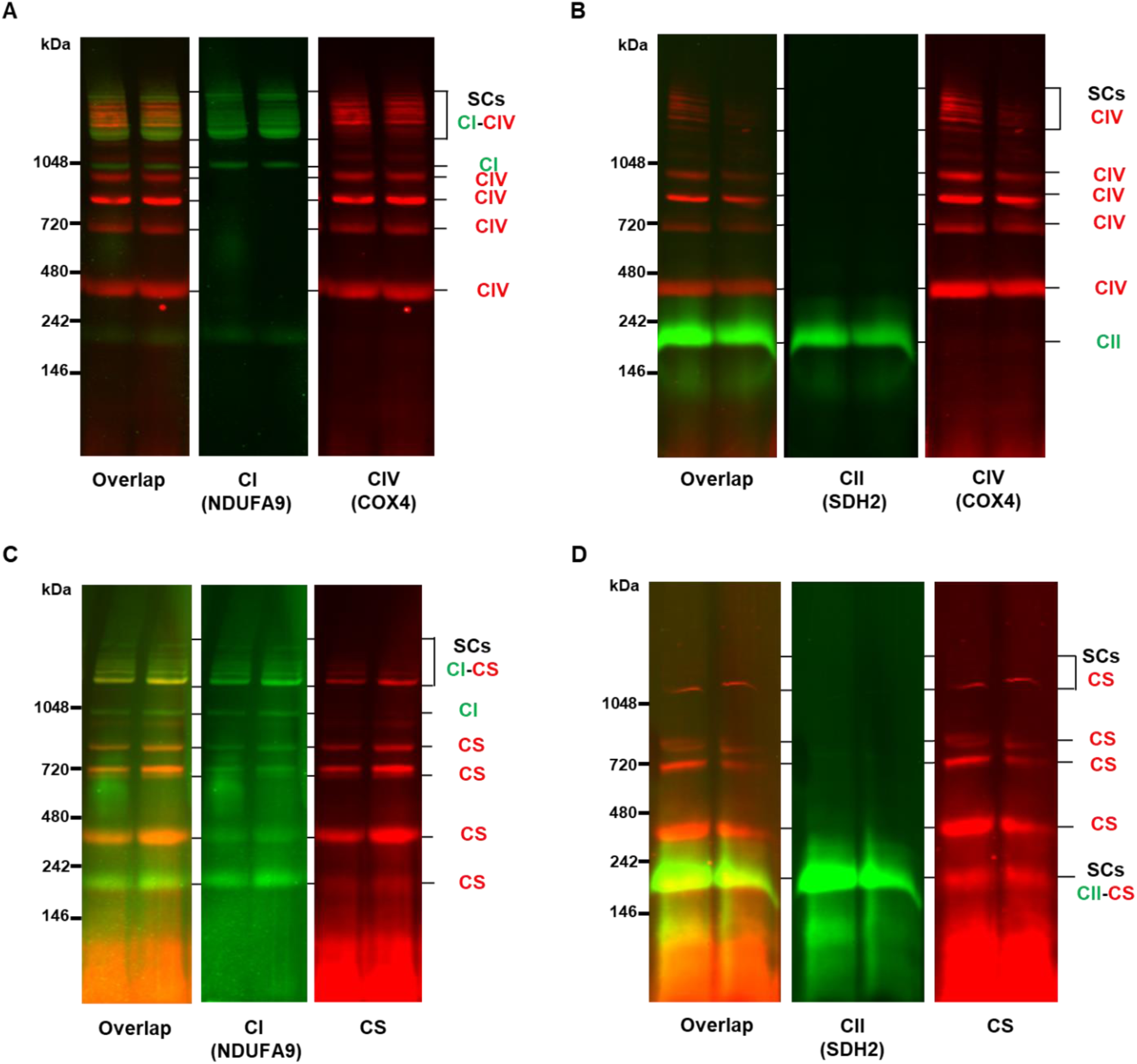
Mitochondrial supercomplexes (SCs) of mouse frozen left ventricular tissue homogenates using blue native gel electrophoresis followed by immunoblotting. (A) Representative images of mitochondrial CI-CIV SCs (colored yellow) and individual bands corresponding to CI (colored green) and CIV (colored red). (B) No CII-CIV SCs were found. Individual bands corresponding to CII (colored green) and CIV (colored red) could be observed. (C) Representative images of SCs formed with CS and CI (colored yellow) and individual bands corresponding to CI (colored green) and CS (colored red). (D) Representative images of SCs formed with CII and CS (colored yellow). Individual bands corresponding to CII (colored green) and CS (colored red). CI: complex I, CII: complex II, CIV: complex IV, CS: citrate synthase.

### The TSIT method reveals significant differences in mitochondrial respiratory function between frozen atria and ventricles in mice

To test if the mitochondrial respiration differences between fresh atrial and ventricular tissues could also be observed in the frozen samples, mitochondrial respiratory capacities of frozen mouse atrial and ventricular tissues were determined with TSIT (*Figure 4A*). As expected, mitochondrial respiratory capacities could be measured in frozen cardiac tissue with multiple injections of substrates or inhibitors of mitochondrial complexes in a single experimental round. Basal respiratory capacity was determined after the injection of malate, NAD^+^ and Cytc. Acetyl-CoA was then injected to drive the utilization of malate and NAD^+^ in the TCA cycle to produce NADH as the substrate for CI, allowing for the measurement of CI-linked mitochondrial respiratory capacity. The third injection with rotenone could fully block the O_2_ flux, proving that the majority of mitochondrial respiration upon addition of acetyl-CoA was CI-linked. The fourth series of injections were the titration with succinate to measure the CII-linked mitochondrial respiratory capacity. The fifth and sixth injection with malonate and antimycin A were to measure Rox for the correction with non-mitochondrial oxygen consumption. Using this method, we observed that acetyl-CoA mediated mitochondrial CI-linked respiratory capacity and succinate driven CII-linked respiratory capacity in frozen ventricular tissue was significantly higher than the corresponding respiratory capacity in the frozen atrial tissue (*Figure 4B* and *C*). This was consistent with data obtained using fresh tissue (*Figure 1C*). Collectively, these data demonstrate that the optimized TSIT using frozen cardiac tissue homogenates can be performed successfully with comparable reliability and sensitivity to the conventional mitochondrial SUIT respiration measurement using fresh tissue regarding CI- and CII-linked mitochondrial respiration.

**Fig. 4:**
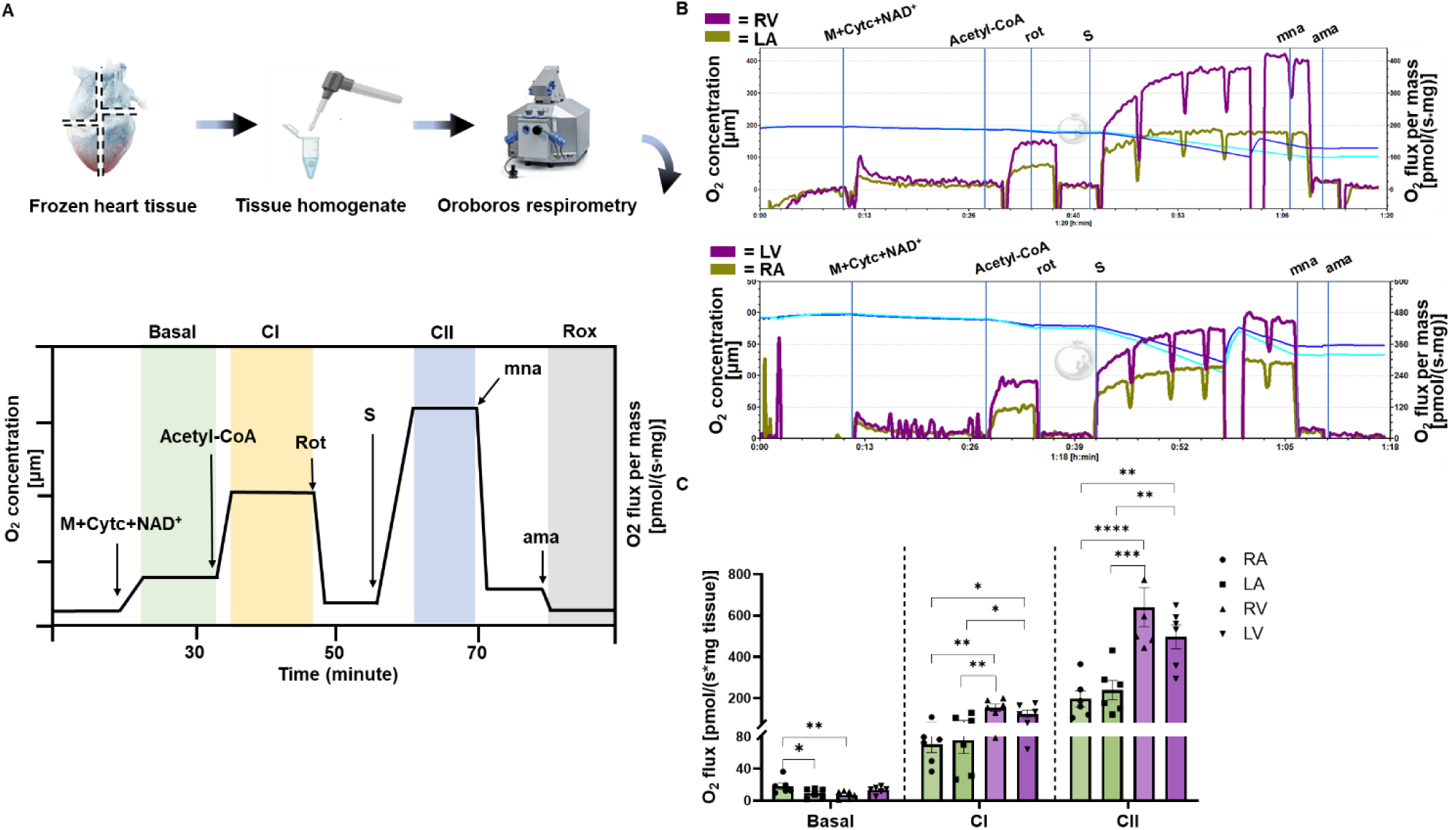
The mitochondrial respiratory capacity of mouse frozen cardiac tissue is comparable to the respiration measured in fresh cardiac tissue. (*A*) Schematic representation of the sample preparation and the TSIT method for frozen cardiac samples from right atrium (RA), left atrium (LA), right ventricle (RV), and left ventricle (LV). Samples were weighed, homogenized and assessed via a high-resolution respirometry using TSIT as illustrated. Basal respiratory capacity (green) was determined at 2 mM malate (M), 100 µM NAD^+^ and 10 µM cytochrome c (Cytc). Complex I (CI)-linked respiratory capacity (yellow) was determined at an additional 150 µM acetyl-CoA. Complex II (CII)-linked respiratory capacity (blue) was determined at 1 µM rotenone (rot) and titration of 20 µl of 1 M succinate (S). Residual oxygen consumption (Rox, gray) was determined at an additional 5 mM malonate (mna) and 5 µM antimycin A (ama). At least 2 technical replicates were performed. (*B*) Representative traces of frozen RA, LA, RV, and LV tissues from BL6JRcc(BL6J)-*Nnt^+^*/Wuhap mice. Purple traces indicate the ventricular O_2_ flux and olive-green traces indicate the atrial O_2_ flux. (*C*) Quantification of basal-, CI- and CII-linked respiratory capacities in frozen atrial and ventricular tissues (n=6). Data are expressed as mean± standard error of the mean. One-way analysis of variance (ANOVA) followed with a post-hoc Fisher’s LSD test was used within each mitochondrial respiratory capacity. \**P* < 0.05, \*\**P* < 0.01, \*\*\**P* < 0.001, \*\*\*\**P* < 0.0001.

### The TSIT method distinguishes differences in mitochondrial respiratory capacity between frozen cardiac tissue of young and old mice

To determine whether TSIT can be applied to distinguish physiologically relevant changes, a well-recognized ageing mouse model was assessed. Using conventional SUIT respirometry protocols, compromised mitochondrial respiration was observed with ageing in mitochondria isolated from fresh mouse LV tissue^40^ and in fresh permeabilized rat ventricular tissue.^41^ Therefore, we performed TSIT respirometry in young (4-month old) and old (23-month old) mice using frozen LV tissue. Our results showed that CII-linked respiration was significantly decreased with age, while the basal and CI-linked respiratory capacities were also decreased with a trend but not reached statistical significance between young and old mice (*Figure 5A* and *B*). These results illustrated TSIT could be applied to detect ageing-related mitochondrial dysfunction in the frozen cardiac tissue.

**Fig. 5:**
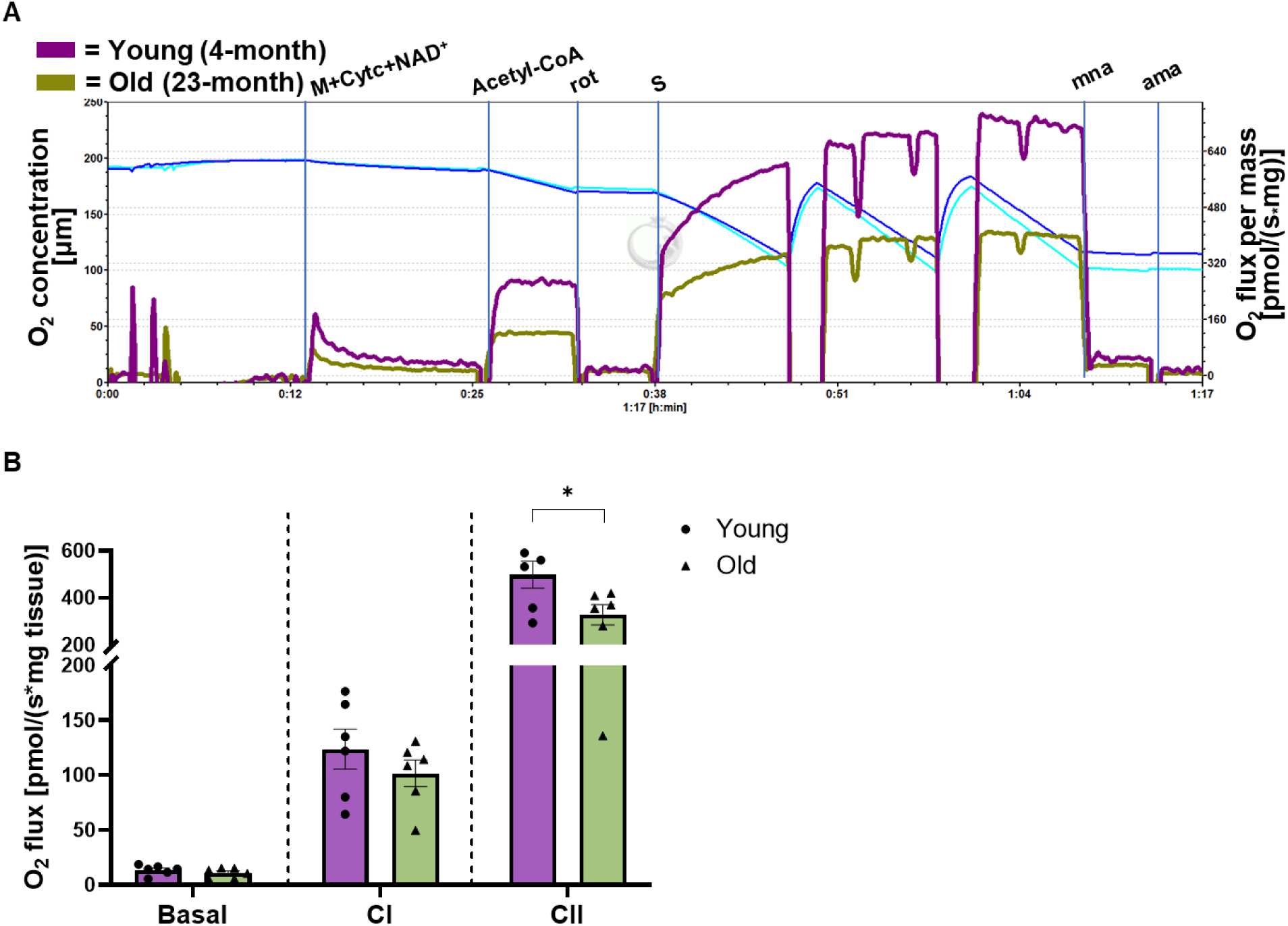
Application of TSIT for high-resolution respirometry of frozen cardiac tissues of young and old mice. (*A*) Representative traces of frozen left ventricular (LV) tissues from young (4-month-old, purple) and old (23-month-old, olive-green) mice. (*B*) Quantification of basal-, complex I (CI)-, and complex II (CII)-linked respiratory capacities in the frozen LV tissues of young (purple) and old (olive-green) mice (n=6). Data are expressed as mean ± standard error of the mean. A two tailed unpaired Student t-test was used within each mitochondrial respiratory capacity. \**P* < 0.05. M+Cytc+NAD^+^: malate+cytochrome c+NAD^+^, rot: rotenone, S: succinate, mna: malonate, ama: antimycin A.

### The TSIT method differentiates the impairment of mitochondrial respiratory capacity in frozen cardiac tissue of pathophysiological disease models

Mitochondrial dysfunction has been associated with a broad range of human diseases. We aimed to apply our TSIT method to measure mitochondrial respiratory (dys)function in frozen cardiac tissue samples from multiple pathophysiological disease models. First, a mouse cardiac IR model was employed, since impaired mitochondrial metabolism, including oxidative stress, mitochondrial calcium overload, and impaired ETC and OXPHOS were reported during cardiac IR injury.^42, 43^ Prolonged ischemia was shown to induce a significant reduction in CI and CII-linked oxygen consumption in rat subsarcolemmal mitochondria isolated from fresh cardiac tissue.^44^ Interestingly, our TSIT method was able to detect a significant impaired CI- and CII-linked respiratory capacities in frozen LV tissue from IR mice compared with the sham control (*Figure 6A* and *B*).

**Fig. 6:**
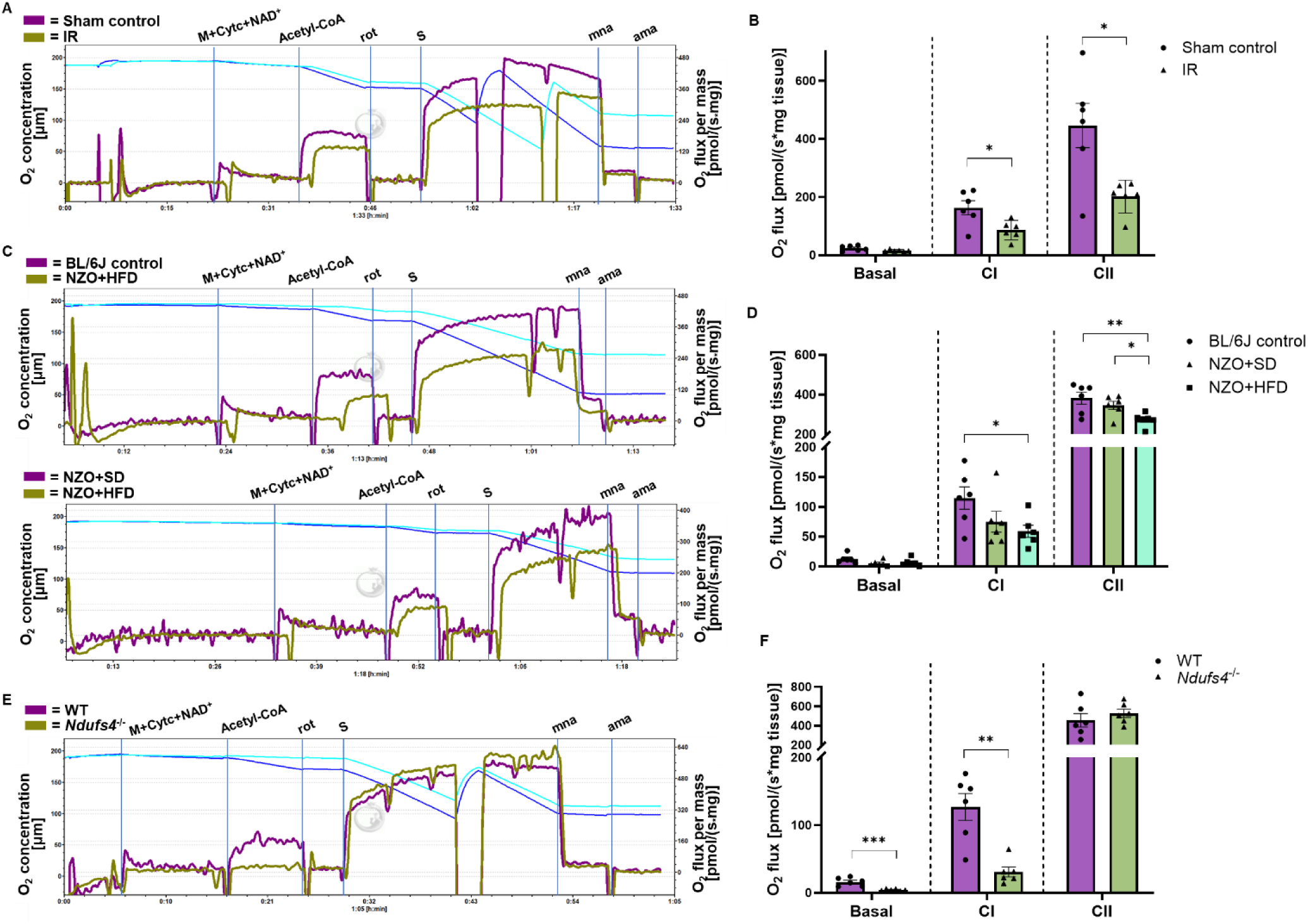
Application of TSIT for high-resolution respirometry of frozen cardiac tissues from pathophysiological disease mouse models. (*A*) Representative traces of frozen left ventricular (LV) tissues from mice subjected to ischemia reperfusion injury (IR; purple) and sham control (olive-green). (*B*) Quantification of basal-, complex I (CI)-, and complex II (CII)-linked respiratory capacities in frozen LV tissues from the IR and the sham control (n=6). (*C*) Representative traces of frozen LV tissues from C57BL/6J mice fed with a standard chow diet (BL/J control; purple), and NZO mice fed with a standard chow diet (NZO+SD; olive-green) or a high fat diet (NZO+HFD; light blue). (*D*) Quantification of basal-, CI-, and CII-linked respiratory capacities in frozen LV tissues from BL/6J, NZO+SD, and NZO+HFD (n=6). (*E*) Representative traces of frozen LV tissues from *Ndufs4* knockout (*Ndufs4*^−/−^; olive-green) and wildtype (WT; purple) mice. (*F*) Quantification of basal-, CI-, and CII-linked respiratory capacities in frozen LV tissues in WT and *Ndufs4*^−/−^ mice (n=6). A two tailed unpaired Student t-test was used within each mitochondrial respiratory capacity. Data are expressed as mean ± standard error of the mean. \**P* < 0.05, \*\**P* < 0.01, \*\*\**P* < 0.001. M+Cytc+NAD^+^: malate+cytochrome c+NAD^+^, rot: rotenone, S: succinate, mna: malonate, ama: antimycin A.

Moreover, we tested TSIT on the NZO mouse model of obesity and type 2 diabetes. Previous research has shown that CII-linked mitochondrial respiration was decreased in fresh LV tissue of diabetic mice.^2^ Similarly, our TSIT method revealed a significant lower CI-linked respiratory capacity in LV tissue of the NZO+HFD group compared with the BL/6J control. Furthermore, CII-linked respiratory capacity was significantly lower in LV tissue of NZO+HFD group compared with both the BL/6J control and NZO+SD groups (*Figure 6C* and *D*).

To further validate TSIT in a genetically modified mouse model with impaired mitochondrial function, we assessed respiratory capacity in frozen cardiac tissue from a CI-deficient (*Ndufs4*^−/−^) mouse model. Reduced activity of mitochondrial CI is expected to significantly impact CI-linked respiration in *Ndufs4*^−/−^ mouse heart. Indeed, a significantly decreased basal and CI-linked respiratory capacity was observed in frozen LV tissue from *Ndufs4*^−/−^ mice compared with the WT control, while CII-linked respiratory capacity remained unchanged (*Figure 6E* and *F*). Taken together, these observations strongly support the conclusion that TSIT is capable of detecting mitochondrial respiratory differences across various pathophysiological conditions using frozen cardiac tissue.

### The TSIT method detects mitochondrial damage in human frozen cardiac tissue

To promote the utility of TSIT in clinical practice, we measured mitochondrial respiratory capacities using frozen heart tissue homogenates from human donors. Five LV samples with cardiac pathology (CP) and four LV samples with no/low cardiac pathology (NCP) were used (Table S1). We could thus measure mitochondrial respiration in frozen human tissue using the same TSIT method with approximately 2 mg of tissue. The TSIT was able to detect a significantly impaired CII-linked respiratory capacity in frozen LV tissue from the CP compared with the NCP, while no significant difference in basal- and CI-linked respiratory capacities between the two groups was observed (*Figure 7A* and *B*).

**Fig. 7:**
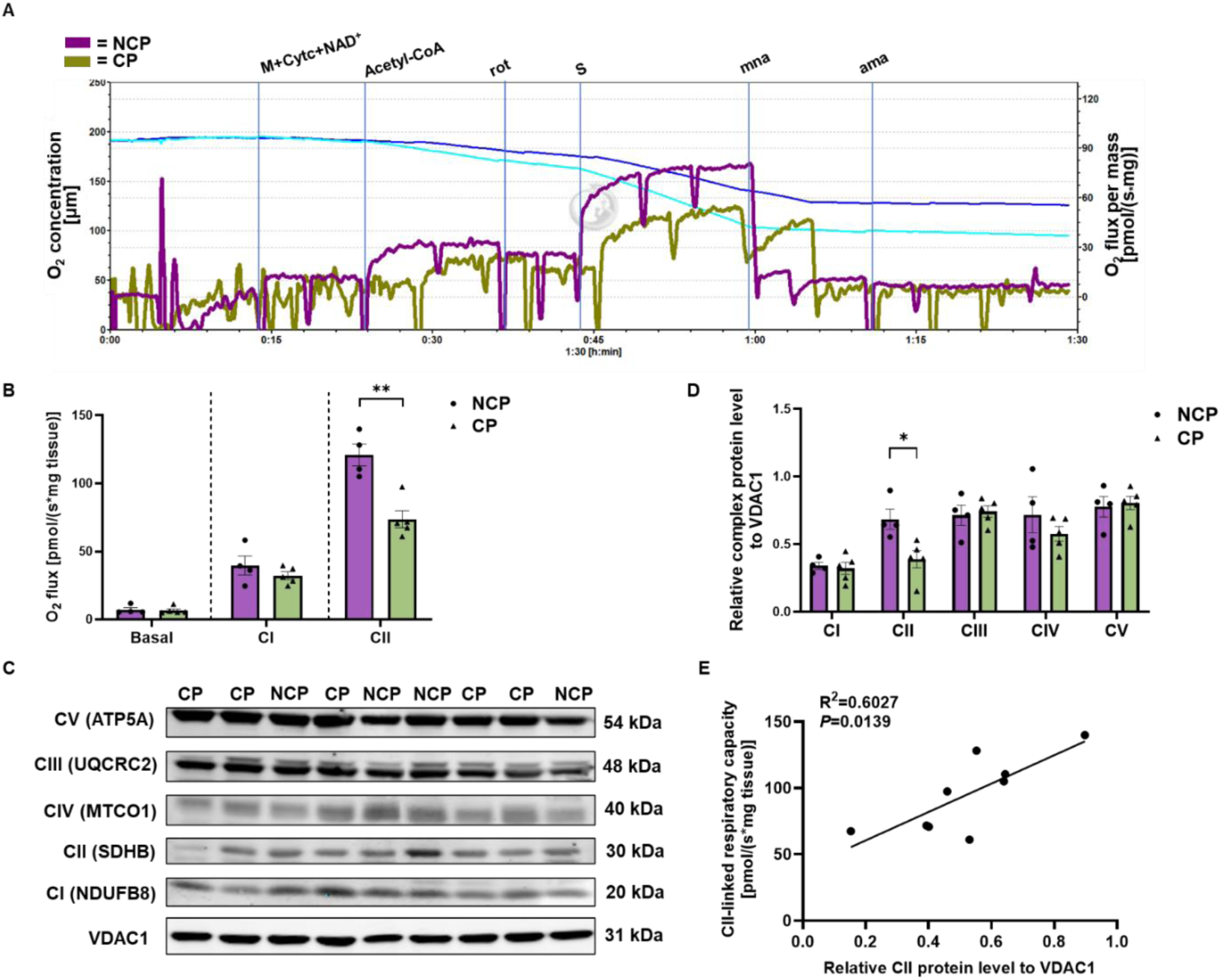
Application of TSIT for high-resolution respirometry of frozen cardiac tissue from human donors with either no/low cardiac pathology or high cardiac pathology. (*A*) Representative traces of frozen left ventricular (LV) tissue from human donors with no/low cardiac pathology (NCP, purple) and with cardiac pathology (CP, olive-green). (*B*) Quantification of basal-, complex I (CI)-, and complex II (CII)-linked respiratory capacities in frozen LV tissues from NCP (n=4) and CP (n=5) groups. A two tailed unpaired Student t-test was used within each mitochondrial respiratory capacity. (C) Representative Western blotting plots of individual mitochondrial CI to CV in frozen LV tissue from NCP and CP groups. VDAC1 protein served as the loading control. (D) Quantification of relative complex protein level of mitochondrial CI to CV to the loading control VDAC1. A two tailed unpaired Student t-test was used within each mitochondrial complex protein level. (E) Pearson correlation analysis between CII-linked respiratory capacity and relative CII protein level to VDAC1 in frozen LV tissue from NCP and CP groups. Data are expressed as mean ± standard error of the mean. \**P* < 0.05, \*\**P* < 0.01. M+Cytc+NAD^+^: malate+cytochrome c+NAD^+^, rot: rotenone, S: succinate, mna: malonate, ama: antimycin A.

To test whether the decreased oxygen consumption in frozen LV tissue measured by TSIT was because of differences in the protein levels of the individual mitochondrial complexes, Western blotting was performed. We observed a significant decrease in CII protein level in the CP group compared with the NCP group, with no difference in the other mitochondrial complexes between the two groups (*Figure 7C* and *D*). Notably, correlation analysis showed that CII-linked respiratory capacity measured by TSIT was significantly correlated with relative CII protein expression (*Figure 7E*). This finding suggests that TSIT is sensitive to identify impairment in specific mitochondrial complexes.

## Discussions

To the best of our knowledge, the current study was the first to establish a method using high-resolution respirometry to assess mitochondrial function of multiple mitochondrial complexes in frozen cardiac tissue with tailored substrate-inhibitor titration (TSIT). TSIT measures oxygen consumption following electron delivery by the TCA cycle as well as by direct CI and CII substrates. The small amount of frozen sample material, a straightforward preparation of tissue homogenates, and unlimited titration with reagents for mitochondrial ETC complexes allow to measure detailed mitochondrial respiratory capacities in frozen cardiac tissue of mice and humans. While for mice 0.4-0.7 mg of atrial and ventricular tissue was used, 1.8-2.2 mg of human ventricular tissue was used. This difference likely reflects the higher metabolic rate of mice compared to humans.^45^ Using cardiac tissues from the same batch of mice, TSIT identified differences in mitochondrial respiration between frozen atrial and ventricular tissues, consistent with those identified using the conventional SUIT protocol with fresh tissue. Moreover, by testing in both physiological and pathophysiological models, TSIT effectively distinguished primary differences or impairment in mitochondrial respiratory function of diseased frozen mouse and human cardiac tissue samples compared with the controls. Therefore, evaluating mitochondrial function in frozen cardiac tissue using the TSIT method could significantly contribute to the laboratory assessment of patients with cardiac diseases.

### Comparison of mitochondrial function between atrial and ventricular tissue

To our knowledge, this is the first study to compare the mitochondrial differences between atrial and ventricular tissues in mice using both conventional SUIT protocol and the newly established TSIT. The differences observed in mitochondrial respiratory capacities between frozen atrial and ventricular tissue were in line with those of Lemieux *et al*. (2011) using fresh tissue from human failure hearts.^46^ They found that respiratory capacities were higher in the ventricles, corresponding to the higher specific CS activity in ventricular compared to atrial tissue. In addition, they also showed that mitochondrial respiration capacities per tissue mass were identical in the right and left atrial tissue, which is similar with what we found in mice.^46^ These concordant results from human and mouse cardiac tissue indicate that the TSIT method can reliably reveal physiologically relevant differences in mitochondrial function.

### Cardiac mitochondrial dysfunction under pathophysiological conditions

Mitochondrial functional evaluation is critical for characterizing the pathophysiological differences and metabolic abnormalities in heart diseases. Our results demonstrate that TSIT can effectively identify mitochondrial ETC defects in frozen cardiac tissue from various mouse (patho)physiological models of cardiac aging, obesity, IR injury, and CI deficiency, all of which are known for exhibiting mitochondrial dysfunction.^47–53^ Notably, utilizing TSIT, we were able to determine an impaired CII-linked respiratory capacity in the frozen human LV tissue with cardiac pathology compared with the control. This finding was further supported by the changes in protein expression of individual mitochondrial CII subunit. These results suggest that TSIT is a sensitive and reliable method for diagnosis of mitochondrial dysfunction in frozen stored human samples in clinical practice. The successful application of TSIT in the evaluation of mitochondrial respiratory function under both physiological and pathophysiological conditions underscores the sensitivity and robustness of the method.

### Supercomplexes formation of mitochondrial complexes and TCA cycle component

The successful assessment of CI- and CII-linked mitochondrial respiratory capacities using TSIT indicated that the ETC SCs, specifically SCs formed by CI and CII to CIV, maintained their associations for electron transport and oxygen consumption. Based on the ‘plasticity model’ of mitochondrial SCs theory,^54^ the mitochondrial complexes can be functional either individually or as part of SCs detected by BNGE. Most CI appeared to form SCs with other complexes including CIII and CIV, or CII, CIII, and CIV.^34, 35^ We indeed observed SCs containing CI and CIV formed in frozen mouse cardiac tissue, possibly enhancing efficiency of electron transport from CI to CIV where oxygen is consumed. Notably, due to the technical limitation of BNGE, no CII-CIV SCs could be detected, although it has been recently proven that CII forms SCs with CI and CIV. ^28, 54, 55^ Inspired by our observation that acetyl-CoA could drive CI-linked mitochondrial respiration, we analyzed whether SCs between CS and ETC complexes existed in the frozen cardiac tissue. This revealed physical interactions between the important TCA cycle enzyme CS with CI as well as CII. To our best knowledge, this is the first study to show that the TCA enzymes form SCs with the ETC complexes. Taken together, in frozen cardiac tissue, not only the mitochondrial ETC-formed SCs, but also the TCA metabolon is preserved and coupled to the ETC SCs. This could also explain why mitochondrial respiratory capacities could be measured by TSIT in frozen cardiac tissue.

### Comparison of different respirometry protocols measuring mitochondrial function in frozen tissue

Although TSIT is capable of recapitulating the mitochondrial respiration measurement as is the traditional SUIT protocol with fresh tissue, the mitochondrial respiratory capacities measured by TSIT should be regarded as the maximal uncoupled respiration, but not OXPHOS-related respiration in the context of conventional respirometry theory using fresh tissue.^20^ This is because the MIM disruption in the freeze-thaw process leads to dissipation of mitochondrial membrane potential and thus full uncoupling, allowing respiration with a maximal flow within the ETC system. In addition, different reagents were required in TSIT compared with the SUIT protocol. For frozen homogenates of human muscle biopsies, an absence of oxygen consumption was seen in response to pyruvate and malate, routinely used reagents in the SUIT protocol designed for fresh tissue.^22, 30^ This impairment is probably linked to the inability to convert pyruvate to acetyl-CoA possibly due to loss of pyruvate dehydrogenase in mitochondrial matrix in frozen tissue. Alternatively, this impairment could be due to the lack of NAD^+^ as substrate for NADH production. In contrast, in TSIT, acetyl-CoA together with sufficient NAD^+^, malate and Cytc were added. Due to the disrupted mitochondrial membranes of frozen tissue, these substrates can feed the TCA cycle to generate NADH. The produced NADH then serves as the direct substrate for CI allowing for the measurement of CI-linked mitochondrial respiratory capacity. This is confirmed by a specifically reduced CI- but not CII-linked respiratory capacity in the *Ndufs4*^−/−^ mice and by the fact of complete inhibition of CI-linked respiratory capacity by rotenone in TSIT. The ability of produced NADH to efficiently boost CI-linked mitochondrial respiratory capacity also serves as a proof for SCs formation between TCA cycle enzymes with CI. Furthermore, a hyper-oxygenated condition during respirometry measurement is normally preferred in the SUIT protocol for fresh permeabilized tissue, while a normal oxygen condition was used in TSIT. This is because such small amount of frozen tissue homogenate was fully homogeneous allowing sufficient oxygen penetration under normal oxygen condition. Moreover, respiratory capacity of frozen tissue homogenate remained consistent in hyper-oxygenated and normal conditions (data not presented), thereby eliminating the potential variance due to differences in oxygen level.

Compared with other respirometry protocols using different types of frozen tissues, TSIT showed advantages from various perspectives. Zuccolotto *et al*. (2021), using frozen skeletal muscle tissue, titrated acetyl-CoA together with NAD^+^, malate and Cytc, and also showed a significant correlation between CI-mediated respiration and enzyme activity of individual CI (r = 0.597).^30^ However, they did not include the measurement of CII-linked mitochondrial respiratory capacity nor did they test the maximum CI capacity driven by NADH as a direct substrate. In TSIT, we are able to measure CII-linked respiratory capacity by titration of succinate, which donates H^+^-linked two-electron to FAD bound to the subunit of CII to form FADH_2_, relaying electrons further via a series of iron-sulfur clusters to ubiquinone (coenzyme Q).^56^ An evident inhibitory effect of malonate confirmed that the succinate is indeed feeding CII in the frozen cardiac tissue in TSIT.^57^ Another recent respirometry protocol using frozen tissue using the Seahorse XF Analyzer was established by Acin-Perez *et al*. (2020).^22^ In the process of sample preparation, they centrifugated the tissue lysate to remove debris. Nevertheless, our study proved that centrifugation of tissue lysate prior to respirometry resulted in the loss of mitochondrial protein and induced bigger technical variation, therefore we recommend omitting centrifugation by using whole tissue homogenate, leading to less technical variation and higher sensitivity. Secondly, the Seahorse XF Analyzer allows for only four predetermined injections of (mixed) reagents, limiting options for titration and verification. The TSIT method is more flexible, illustrated by the inclusion of a separate injection of rotenone to confirm measurement of CI-linked respiratory capacity. Lastly, they used NADH to assess CI-linked respiration, given their hypothesis of an impaired TCA cycle in frozen tissue.^22^ However, their proposition is challenged by our data and data from Zuccolotto *et al*.,^30^ showing that the TCA cycle remained functional in frozen cardiac and muscle tissue as NADH could be produced by TCA cycle with the supply of acetyl-CoA, NAD^+^, malate and Cytc. This is also supported by SCs formation of ETC SCs with the TCA cycle metabolon. Furthermore, the respiration assessed by direct titration of NADH determines the maximum capacity of CI utilizing the substrate beyond its availability in the physiological condition, which is mostly produced via the TCA cycle. Therefore, the CI-linked respiration fed by NADH produced from TCA cycle could better represent TCA cycle-coupled respiration process within mitochondria.

### Limitations

Unfortunately, TSIT could not overcome some limitations of using frozen tissue. A limitation of our method is its inability to assess the FAO-linked respiration even with the necessary co-factors for FAO. This is probably due to the disassembly, inactivation or disintegration of FAO-related enzymatic complexes preventing them from donating electrons to the coenzyme Q or providing acetyl-CoA to the TCA cycle. It is also important to point out that the interference from varying times of freezing or from sampling of different areas, especially in large human cardiac tissue, cannot be excluded. However, the same limitation exists in conventional respirometry protocols using fresh tissue. These effects need to be studied, in order to provide solid frozen tissue biobanking advise. Furthermore, we showed the potential association of metabolon association with ETC SCs with a single method. This opens a new avenue of research. However, future studies are warranted to further confirm this conclusion.

In conclusion, the newly developed versatile TSIT using high resolution respirometry was shown to be a valid and robust method to evaluate mitochondrial respiratory capacities in diverse frozen cardiac tissues from different species, distinguishing physiological and pathophysiological impairments, enhancing mitochondrial function analyses for both laboratory and clinical applications.

## Acknowledgements

We appreciate the technical assistance from Soumi Ganguli, Haomiao Wang, Rijk de Jong, and Daan Hanegraaf from Human and Animal Physiology, and Jelmer Vroom from the Laboratory of Virology.

## Ethics declarations

The authors declare no competing interests.

## Authors’ contributions

D.Z. obtained the funding and launched the study. L.H. and D.Z. conceived the ideas and designed the experiments. L.H., S.F., and X.J. carried out the experimental work, analyzed the data, and wrote the manuscript draft. A.R., J.K., and D.Z. provided scientific feedback on experiments and revised the manuscript. R.C., M.G., C.O., M.W., and W.K. provided sample materials for testing the application of the method and provided critical feedback. J.M. M.O., and M.B. provided technical assistance and technical training. All the authors read and commented on the manuscript.

## Supplementary

**Table S1.**
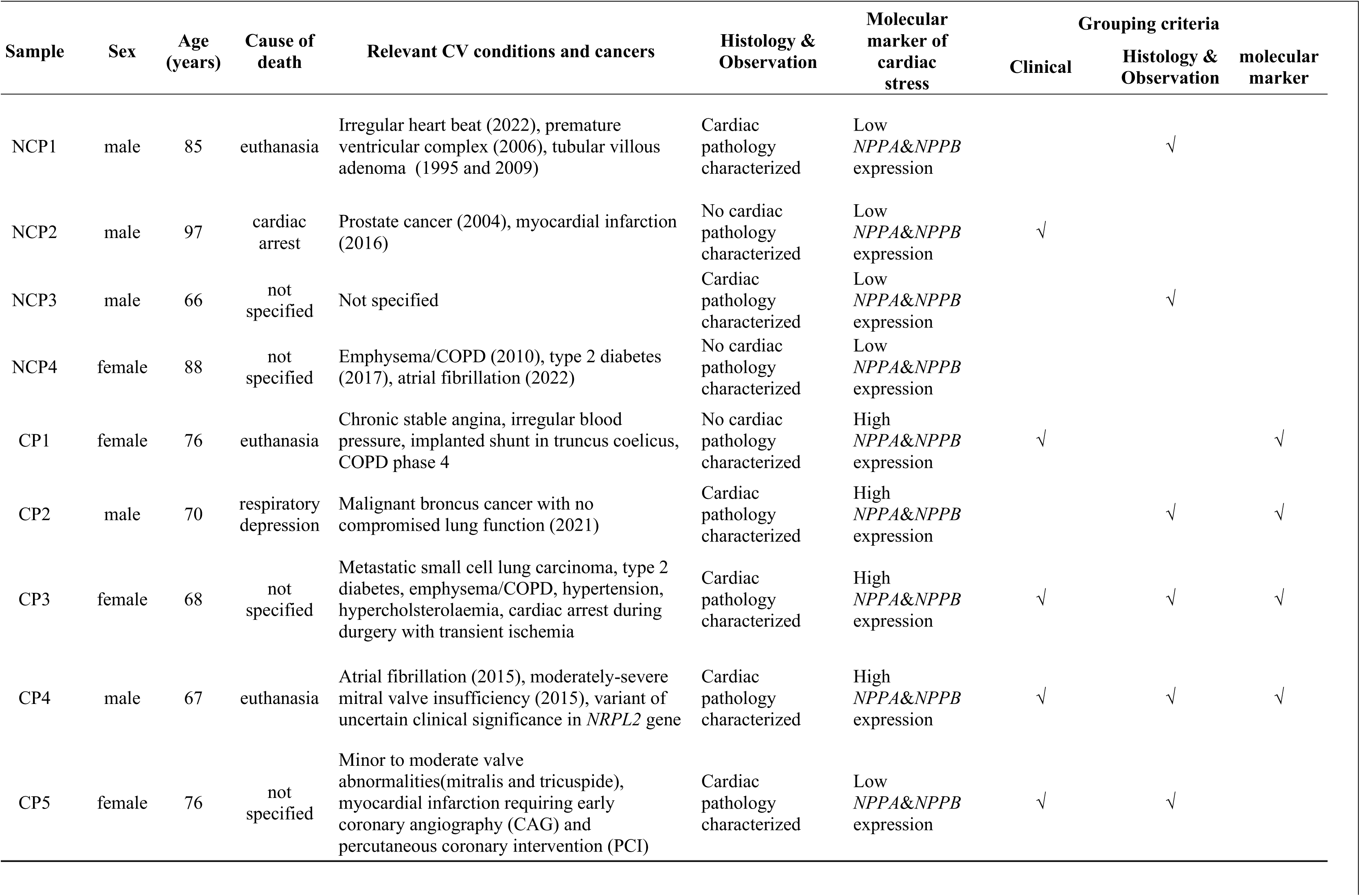
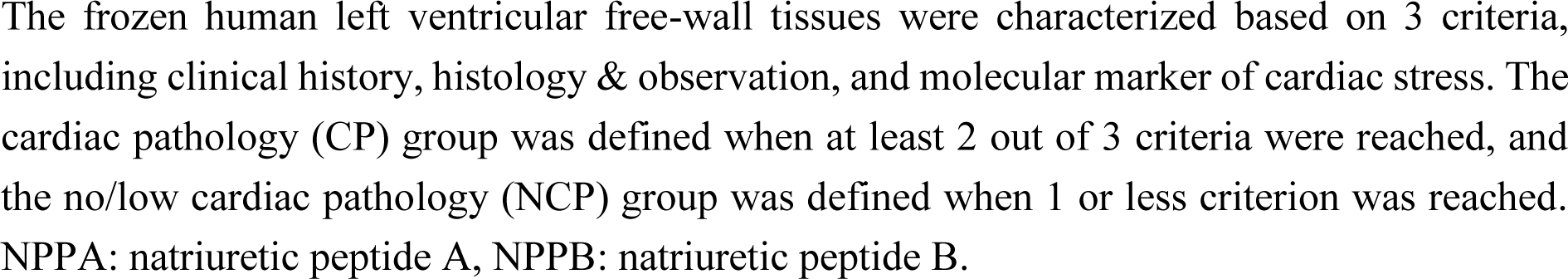
Basic clinical & molecular information of human donors.

**Fig. S1:**
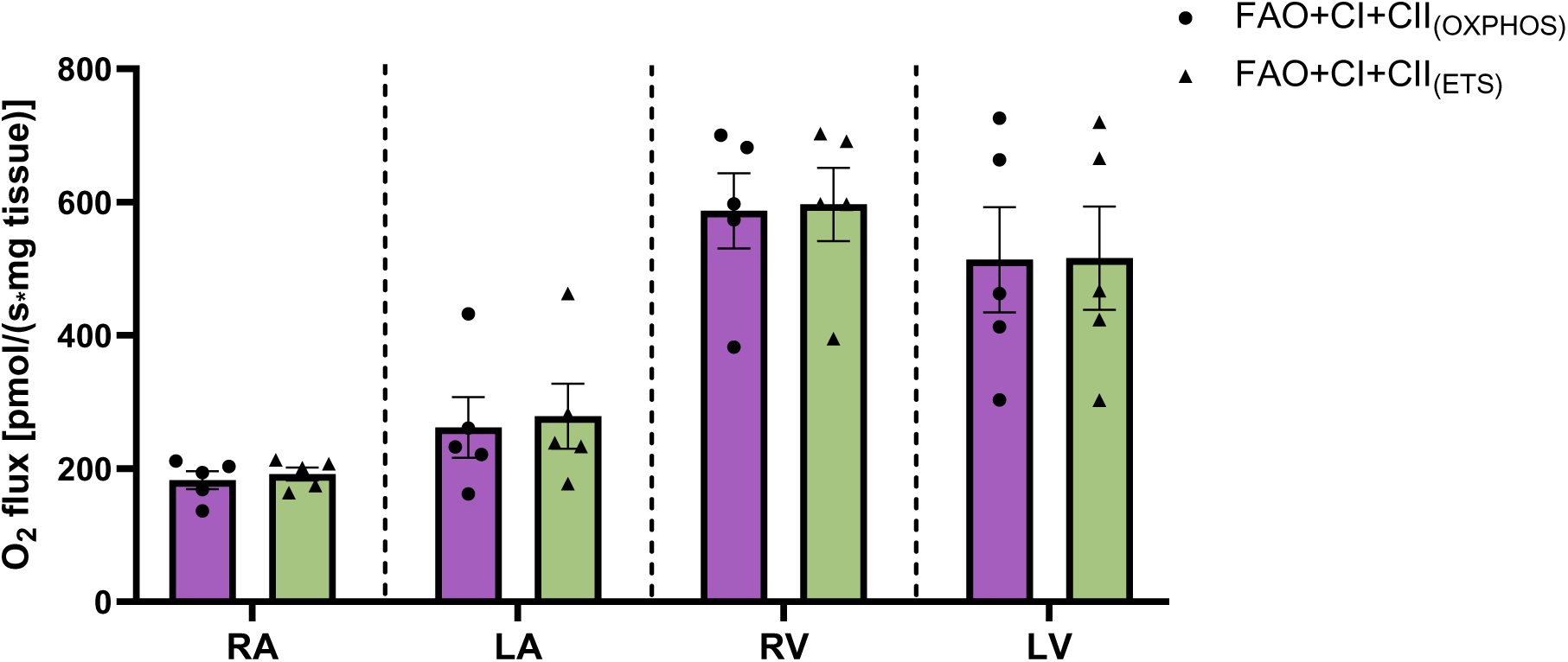
Mitochondrial respiration of the oxidative phosphorylation (OXPHOS) state and the uncoupled maximal electron transport chain capacity (ETS state) in fresh mouse cardiac tissues. The fatty acid oxidation+complex I+complex II (FAO+CI+CII)-linked mitochondrial respiration in the OXPHOS state (purple) was determined at 7.5 mM ADP, 0.3 mM malate, 0.5 mM octanoylcarnitine, 2 mM malate, 5 mM pyruvate, 10 mM glutamate, and 10 mM succinate. FAO+CI+CII-linked mitochondrial respiration in the ETS state (olive-green) was determined at an additional titration of 1 μl of 0.1 μM carbonyl cyanide-p-trifluoromethoxyphenylhydrazone (CCCP). No differences were observed between these two states (n=5). Data are expressed as mean ± standard error of the mean. Individual group mean differences within each cardiac tissue type were tested with unpaired t test. RA: right atrium, LA: left atrium, RV: right ventricle, LV: left ventricle.

**Fig. S2:**
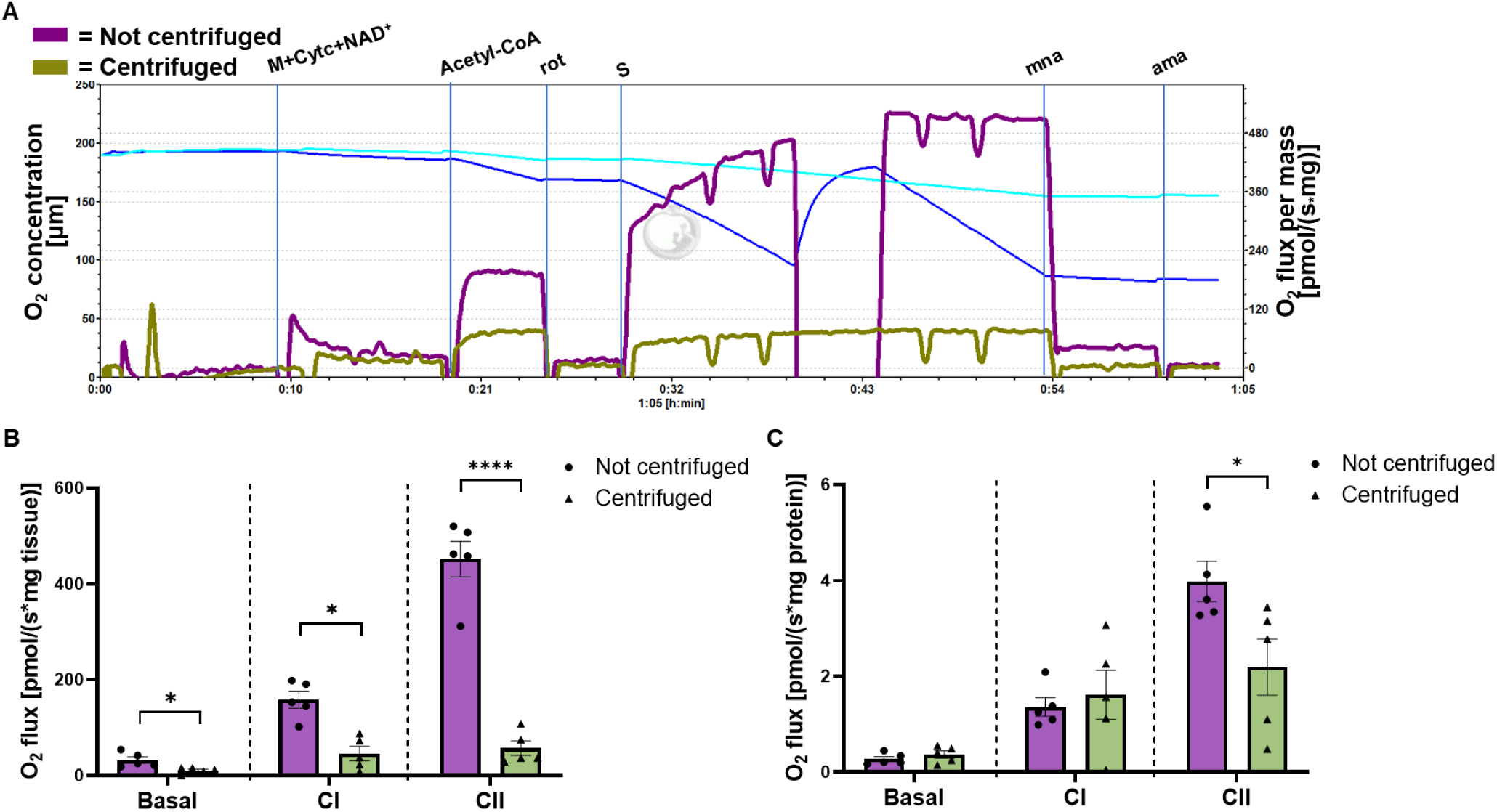
Influence of centrifugation of frozen mouse cardiac tissue homogenates on respirometry measurement using the TSIT method. (A) Representative trace of high-resolution respirometry of frozen left ventricular tissue homogenates with (olive-green) and without centrifugation (purple). (B) Quantified O_2_ flux showed that basal, complex I (CI)- and complex II (CII)-linked respiratory capacities were all significantly hampered by the centrifugation at the speed of 1000 x *g* for 5 min at 4 °C prior to the respirometry measurement (n=5). (C) Corrected O_2_ flux to protein concentration revealed that the significant differences between centrifuged and uncentrifuged samples were less pronounced, primarily affecting the data obtained from the centrifuged samples. Data are expressed as mean ± standard error of the mean. A two tailed unpaired Student t-test was used within each mitochondrial respiratory capacity. \**P* < 0.05, \*\*\**P* < 0.001, \*\*\*\**P* < 0.0001. M+Cytc+NAD^+^: malate+cytochrome c+NAD^+^, rot: rotenone, S: succinate, mna: malonate, ama: antimycin A.

**Fig. S3:**
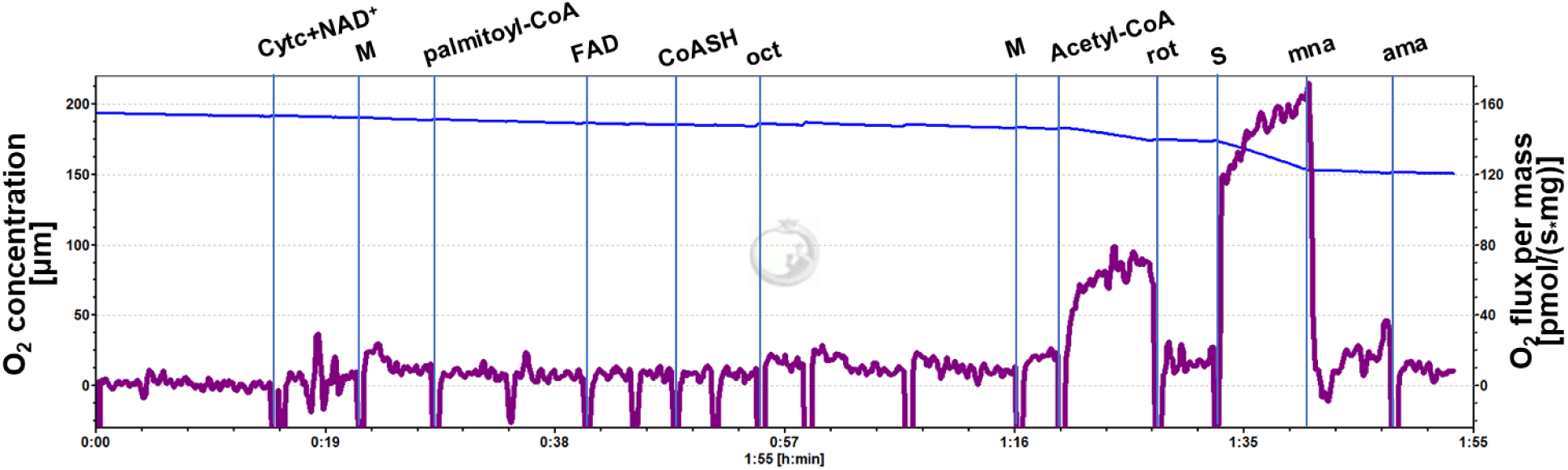
Representative trace of high-resolution respirometry for the measurement of fatty acid oxidation (FAO) in frozen mouse left ventricular tissues. Basal respiratory capacity was determined at 100 µM NAD^+^,10 µM cytochrome c (Cytc) and 2 mM malate (M). FAO-linked respiratory capacity was determined at an additional titration of 40 µM palmitoyl-CoA, 40 µM FAD, and 1.2mM coenzyme A (CoASH). FAD is necessary for first step of FAO when the acyl CoA (e.g. palmityl CoA) is oxidized to an enoyl CoA molecule by acyl CoA dehydrogenase. Palmityl-CoA as the substrate of FAO, instead of palmityl-carnitine, was added because the long chain fatty acyl-CoA can cross the broken mitochondrial membrane due to the freeze-thaw process. CoASH is the necessary co-factor in the final step of FAO. However, the flux remained unchanged. Also no O_2_ flux was observed at an additional titration of 0.5 mM octanoylcarnitine (oct), but a substantial flux after addition of 150 µM acetyl-CoA (CI-linked respiratory capacity) and 10 mM succinate (S, CII-linked respiratory capacity) indicated that the TCA cycle, CI and CII were still functional and linked. CI: complex I, CII: complex II, rot: rotenone, mna: malonate, ama: antimycin A.

**Fig. S4:**
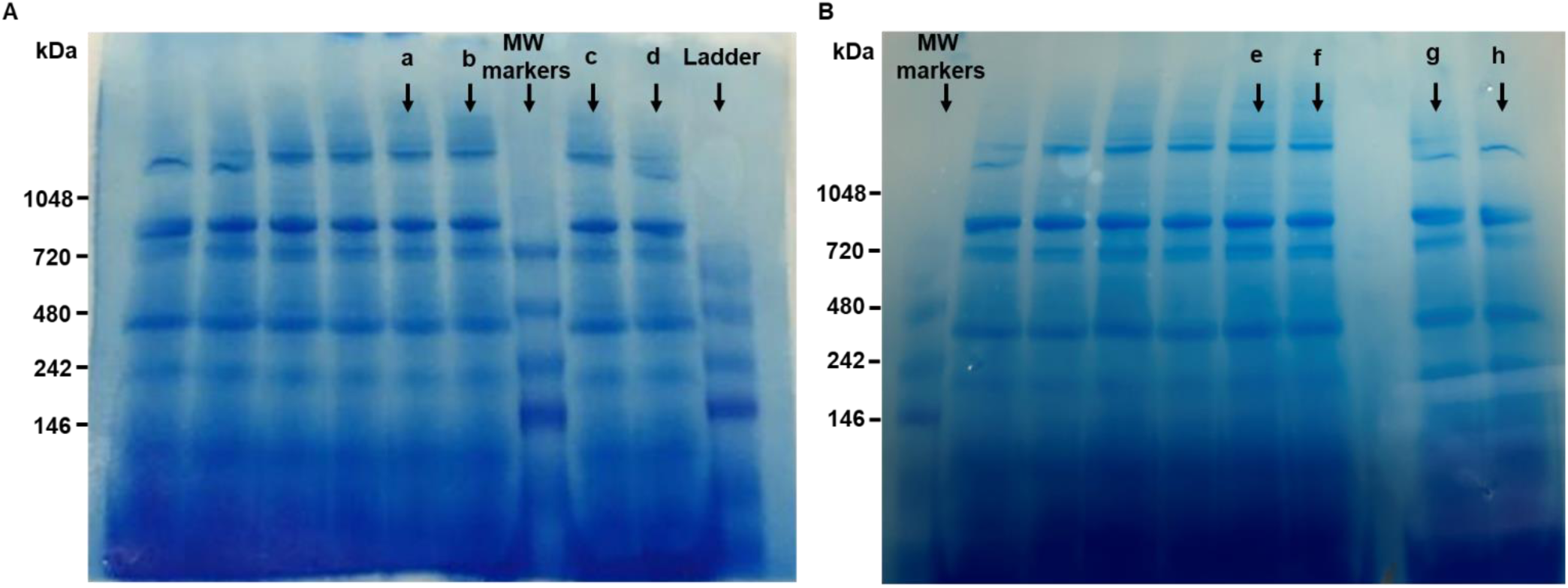
Loading control of blue native gel electrophoresis stained with coomassie brilliant blue G. (A) Blue native gel for the measurement of supercomplexes (SCs) between CI, CII and CIV in mouse frozen left ventricular (LV) tissue homogenates. Lane ‘a’ and ‘b’ were the representative blots of SCs CI-CIV; ‘c’ and ‘d’ were the representative blots of SCs CII-CIV. (B) Blue native gel for the measurement of SCs between CI, CII and CS in mouse frozen LV tissue homogenates. Lane ‘e’ and ‘f’ were the representative blots of SCs CI-CS; ‘g’ and ‘h’ were the representative blots of SCs CII-CS. CI: complex I, CII: complex II, CIV: complex IV, CS: citrate synthase.

## References

1. Müller MJ, Wang Z, Heymsfield SB, Schautz B, Bosy-Westphal A. Advances in the understanding of specific metabolic rates of major organs and tissues in humans. Curr Opin Clin Nutr Metab Care 2013;16:501–508.

2. Parker AM, Tate M, Prakoso D, Deo M, Willis AM, Nash DM, Donner DG, Crawford S, Kiriazis H, Granata C, Coughlan MT, De Blasio MJ, Ritchie RH. Characterisation of the Myocardial Mitochondria Structural and Functional Phenotype in a Murine Model of Diabetic Cardiomyopathy. Front Physiol 2021;12:672252.

3. Hoeks J, Hesselink M, Schrauwen P. Mitochondrial Respiration. In: Mooren FC, ed. Encyclopedia of Exercise Medicine in Health and Disease. Berlin, Heidelberg: Springer Berlin Heidelberg, 2012:587–590.

4. Kumar AA, Kelly DP, Chirinos JA. Mitochondrial Dysfunction in Heart Failure With Preserved Ejection Fraction. Circulation 2019;139:1435–1450.

5. Nollet EE, Duursma I, Rozenbaum A, Eggelbusch M, Wüst RCI, Schoonvelde SAC, Michels M, Jansen M, van der Wel NN, Bedi KC, Margulies KB, Nirschl J, Kuster DWD, van der Velden J. Mitochondrial dysfunction in human hypertrophic cardiomyopathy is linked to cardiomyocyte architecture disruption and corrected by improving NADH-driven mitochondrial respiration. Eur Heart J 2023;44:1170–1185.

6. Walters AM, Porter GA, Brookes PS. Mitochondria as a Drug Target in Ischemic Heart Disease and Cardiomyopathy. Circulation Research 2012;111:1222–1236.

7. Tatarková Z, Kuka S, Račay P, Lehotský J, Dobrota D, Mištuna D, Kaplán P. Effects of aging on activities of mitochondrial electron transport chain complexes and oxidative damage in rat heart. Physiol Res 2011;60:281–289.

8. Tocchi A, Quarles EK, Basisty N, Gitari L, Rabinovitch PS. Mitochondrial dysfunction in cardiac aging. Biochim Biophys Acta 2015;1847:1424–1433.

9. Karamanlidis G, Lee CF, Garcia-Menendez L, Kolwicz SC, Jr., Suthammarak W, Gong G, Sedensky MM, Morgan PG, Wang W, Tian R. Mitochondrial complex I deficiency increases protein acetylation and accelerates heart failure. Cell Metab 2013;18:239–250.

10. Bolea I, Gella A, Sanz E, Prada-Dacasa P, Menardy F, Bard AM, Machuca-Márquez P, Eraso-Pichot A, Mòdol-Caballero G, Navarro X, Kalume F, Quintana A. Defined neuronal populations drive fatal phenotype in a mouse model of Leigh syndrome. eLife 2019;8:e47163.

11. Scheubel RJ, Tostlebe M, Simm A, Rohrbach S, Prondzinsky R, Gellerich FN, Silber R-E, Holtz J. Dysfunction of mitochondrial respiratory chain complex I in human failing myocardium is not due to disturbed mitochondrial gene expression. Journal of the American College of Cardiology 2002;40:2174–2181.

12. Sharov VG, Todor AV, Silverman N, Goldstein S, Sabbah HN. Abnormal mitochondrial respiration in failed human myocardium. Journal of molecular and cellular cardiology 2000;32:2361–2367.

13. Lemieux H, Semsroth S, Antretter H, Höfer D, Gnaiger E. Mitochondrial respiratory control and early defects of oxidative phosphorylation in the failing human heart. Int J Biochem Cell Biol 2011;43:1729–1738.

14. Chen Y, Liu Y, Dorn GW, 2nd. Mitochondrial fusion is essential for organelle function and cardiac homeostasis. Circ Res 2011;109:1327–1331.

15. Croston TL, Thapa D, Holden AA, Tveter KJ, Lewis SE, Shepherd DL, Nichols CE, Long DM, Olfert IM, Jagannathan R, Hollander JM. Functional deficiencies of subsarcolemmal mitochondria in the type 2 diabetic human heart. Am J Physiol Heart Circ Physiol 2014;307:H54–65.

16. Montaigne D, Marechal X, Coisne A, Debry N, Modine T, Fayad G, Potelle C, Arid J-ME, Mouton S, Sebti Y, Duez H, Preau S, Remy-Jouet I, Zerimech F, Koussa M, Richard V, Neviere R, Edme J-L, Lefebvre P, Staels B. Myocardial Contractile Dysfunction Is Associated With Impaired Mitochondrial Function and Dynamics in Type 2 Diabetic but Not in Obese Patients. Circulation 2014;130:554–564.

17. Anderson EJ, Kypson AP, Rodriguez E, Anderson CA, Lehr EJ, Neufer PD. Substrate-specific derangements in mitochondrial metabolism and redox balance in the atrium of the type 2 diabetic human heart. J Am Coll Cardiol 2009;54:1891–1898.

18. John C, Grune J, Ott C, Nowotny K, Deubel S, Kühne A, Schubert C, Kintscher U, Regitz-Zagrosek V, Grune T. Sex Differences in Cardiac Mitochondria in the New Zealand Obese Mouse. Front Endocrinol (Lausanne) 2018;9:732.

19. Paradies G, Petrosillo G, Pistolese M, Ruggiero FM. The effect of reactive oxygen species generated from the mitochondrial electron transport chain on the cytochrome c oxidase activity and on the cardiolipin content in bovine heart submitochondrial particles. FEBS Letters 2000;466:323–326.

20. Gnaiger E. Mitochondrial pathways and respiratory control: an introduction to OXPHOS analysis. Bioenergetics communications 2020;2020:2–2.

21. Skladal D, Sperl W, Schranzhofer R, Krismer M, Gnaiger E. Preservation of mitochondrial functions in human skeletal muscle during storage in high energy preservation solution (HEPS). In: What is Controlling Life? Modern Trends in BioThermoKinetics 3 1994:268–271.

22. Acin-Perez R, Benador IY, Petcherski A, Veliova M, Benavides GA, Lagarrigue S, Caudal A, Vergnes L, Murphy AN, Karamanlidis G, Tian R, Reue K, Wanagat J, Sacks H, Amati F, Darley-Usmar VM, Liesa M, Divakaruni AS, Stiles L, Shirihai OS. A novel approach to measure mitochondrial respiration in frozen biological samples. Embo j 2020;39:e104073.

23. Kuznetsov AV, Kunz WS, Saks V, Usson Y, Mazat J-P, Letellier T, Gellerich FN, Margreiter R. Cryopreservation of mitochondria and mitochondrial function in cardiac and skeletal muscle fibers. Analytical Biochemistry 2003;319:296–303.

24. Yamaguchi R, Andreyev A, Murphy AN, Perkins GA, Ellisman MH, Newmeyer DD. Mitochondria frozen with trehalose retain a number of biological functions and preserve outer membrane integrity. Cell Death Differ 2007;14:616–624.

25. Larsen S, Wright-Paradis C, Gnaiger E, Helge JW, Boushel R. Cryopreservation of human skeletal muscle impairs mitochondrial function. Cryo Letters 2012;33:170–176.

26. García-Roche M, Casal A, Carriquiry M, Radi R, Quijano C, Cassina A. Respiratory analysis of coupled mitochondria in cryopreserved liver biopsies. Redox Biology 2018;17:207–212.

27. Barrientos A, Fontanesi F, Díaz F. Evaluation of the mitochondrial respiratory chain and oxidative phosphorylation system using polarography and spectrophotometric enzyme assays. Curr Protoc Hum Genet 2009;Chapter 19:Unit19.13.

28. Mühleip A, Flygaard RK, Baradaran R, Haapanen O, Gruhl T, Tobiasson V, Maréchal A, Sharma V, Amunts A. Structural basis of mitochondrial membrane bending by the I–II–III2–IV2 supercomplex. Nature 2023;615:934–938.

29. Jang DH, Greenwood JC, Spyres MB, Eckmann DM. Measurement of Mitochondrial Respiration and Motility in Acute Care: Sepsis, Trauma, and Poisoning. J Intensive Care Med 2017;32:86–94.

30. Zuccolotto-Dos-Reis FH, Escarso SHA, Araujo JS, Espreafico EM, Alberici LC, Sobreira C. Acetyl-CoA-driven respiration in frozen muscle contributes to the diagnosis of mitochondrial disease. Eur J Clin Invest 2021;51:e13574.

31. Gladka MM, Kohela A, Molenaar B, Versteeg D, Kooijman L, Monshouwer-Kloots J, Kremer V, Vos HR, Huibers MMH, Haigh JJ, Huylebroeck D, Boon RA, Giacca M, van Rooij E. Cardiomyocytes stimulate angiogenesis after ischemic injury in a ZEB2-dependent manner. Nat Commun 2021;12:84.

32. Calvaruso MA, Willems P, van den Brand M, Valsecchi F, Kruse S, Palmiter R, Smeitink J, Nijtmans L. Mitochondrial complex III stabilizes complex I in the absence of NDUFS4 to provide partial activity. Human Molecular Genetics 2011;21:115–120.

33. Costanzini A, Sgarbi G, Maresca A, Del Dotto V, Solaini G, Baracca A. Mitochondrial Mass Assessment in a Selected Cell Line under Different Metabolic Conditions. Cells 2019;8.

34. Enríquez JA. Supramolecular Organization of Respiratory Complexes. Annu Rev Physiol 2016;78:533–561.

35. Lenaz G, Genova ML. Supramolecular organisation of the mitochondrial respiratory chain: a new challenge for the mechanism and control of oxidative phosphorylation. Adv Exp Med Biol 2012;748:107–144.

36. Novack GV, Galeano P, Castaño EM, Morelli L. Mitochondrial Supercomplexes: Physiological Organization and Dysregulation in Age-Related Neurodegenerative Disorders. Front Endocrinol (Lausanne) 2020;11:600.

37. Wu F, Minteer S. Krebs Cycle Metabolon: Structural Evidence of Substrate Channeling Revealed by Cross-Linking and Mass Spectrometry. Angewandte Chemie International Edition 2015;54:1851–1854.

38. Bulutoglu B, Garcia KE, Wu F, Minteer SD, Banta S. Direct Evidence for Metabolon Formation and Substrate Channeling in Recombinant TCA Cycle Enzymes. ACS Chemical Biology 2016;11:2847–2853.

39. Xu X, Liu Y, Luan J, Liu R, Wang Y, Liu Y, Xu A, Zhou B, Han F, Shang W. Effect of downregulated citrate synthase on oxidative phosphorylation signaling pathway in HEI-OC1 cells. Proteome Sci 2022;20:14.

40. Eisenberg T, Abdellatif M, Schroeder S, Primessnig U, Stekovic S, Pendl T, Harger A, Schipke J, Zimmermann A, Schmidt A, Tong M, Ruckenstuhl C, Dammbrueck C, Gross AS, Herbst V, Magnes C, Trausinger G, Narath S, Meinitzer A, Hu Z, Kirsch A, Eller K, Carmona-Gutierrez D, Büttner S, Pietrocola F, Knittelfelder O, Schrepfer E, Rockenfeller P, Simonini C, Rahn A, Horsch M, Moreth K, Beckers J, Fuchs H, Gailus-Durner V, Neff F, Janik D, Rathkolb B, Rozman J, de Angelis MH, Moustafa T, Haemmerle G, Mayr M, Willeit P, von Frieling-Salewsky M, Pieske B, Scorrano L, Pieber T, Pechlaner R, Willeit J, Sigrist SJ, Linke WA, Mühlfeld C, Sadoshima J, Dengjel J, Kiechl S, Kroemer G, Sedej S, Madeo F. Cardioprotection and lifespan extension by the natural polyamine spermidine. Nat Med 2016;22:1428–1438.

41. Jedlička J, Tůma Z, Razak K, Kunc R, Kala A, Proskauer Pena S, Lerchner T, Ježek K, Kuncová J. Impact of aging on mitochondrial respiration in various organs. Physiol Res 2022;71:S227–s236.

42. Kuznetsov AV, Javadov S, Margreiter R, Grimm M, Hagenbuchner J, Ausserlechner MJ. The Role of Mitochondria in the Mechanisms of Cardiac Ischemia-Reperfusion Injury. Antioxidants 2019;8:454.

43. Marin W, Marin D, Ao X, Liu Y. Mitochondria as a therapeutic target for cardiac ischemia-reperfusion injury (Review). Int J Mol Med 2021;47:485–499.

44. Leistner M, Sommer S, Kanofsky P, Leyh R, Sommer S-P. Ischemia time impacts on respiratory chain functions and Ca2+-handling of cardiac subsarcolemmal mitochondria subjected to ischemia reperfusion injury. Journal of Cardiothoracic Surgery 2019;14:92.

45. Perlman RL. Mouse models of human disease: An evolutionary perspective. Evolution, Medicine, and Public Health 2016;2016:170–176.

46. Lemieux H, Semsroth S, Antretter H, Höfer D, Gnaiger E. Mitochondrial respiratory control and early defects of oxidative phosphorylation in the failing human heart. The International Journal of Biochemistry & Cell Biology 2011;43:1729–1738.

47. Duicu OM, Mirica SN, Gheorgheosu DE, Privistirescu AI, Fira-Mladinescu O, Muntean DM. Ageing-induced decrease in cardiac mitochondrial function in healthy rats. Canadian Journal of Physiology and Pharmacology 2013;91:593–600.

48. Lemieux H, Vazquez EJ, Fujioka H, Hoppel CL. Decrease in mitochondrial function in rat cardiac permeabilized fibers correlates with the aging phenotype. J Gerontol A Biol Sci Med Sci 2010;65:1157–1164.

49. Boardman NT, Pedersen TM, Rossvoll L, Hafstad AD, Aasum E. Diet-induced obese mouse hearts tolerate an acute high-fatty acid exposure that also increases ischemic tolerance. Am J Physiol Heart Circ Physiol 2020;319:H682–h693.

50. Guarini G, Kiyooka T, Ohanyan V, Pung YF, Marzilli M, Chen YR, Chen CL, Kang PT, Hardwick JP, Kolz CL, Yin L, Wilson GL, Shokolenko I, Dobson JG, Jr., Fenton R, Chilian WM. Impaired coronary metabolic dilation in the metabolic syndrome is linked to mitochondrial dysfunction and mitochondrial DNA damage. Basic Res Cardiol 2016;111:29.

51. Tompkins AJ, Burwell LS, Digerness SB, Zaragoza C, Holman WL, Brookes PS. Mitochondrial dysfunction in cardiac ischemia–reperfusion injury: ROS from complex I, without inhibition. Biochimica et Biophysica Acta (BBA) - Molecular Basis of Disease 2006;1762:223–231.

52. Kruse SE, Watt WC, Marcinek DJ, Kapur RP, Schenkman KA, Palmiter RD. Mice with Mitochondrial Complex I Deficiency Develop a Fatal Encephalomyopathy. Cell Metabolism 2008;7:312–320.

53. Quintana A, Kruse SE, Kapur RP, Sanz E, Palmiter RD. Complex I deficiency due to loss of Ndufs4 in the brain results in progressive encephalopathy resembling Leigh syndrome. Proceedings of the National Academy of Sciences 2010;107:10996–11001.

54. Acín-Pérez R, Fernández-Silva P, Peleato ML, Pérez-Martos A, Enriquez JA. Respiratory active mitochondrial supercomplexes. Mol Cell 2008;32:529–539.

55. Schon EA, Dencher NA. Heavy breathing: energy conversion by mitochondrial respiratory supercomplexes. Cell Metab 2009;9:1–3.

56. Gnaiger E. Complex II ambiguities—FADH2 in the electron transfer system. Journal of Biological Chemistry 2024;300.

57. Wojtovich AP, Brookes PS. The endogenous mitochondrial complex II inhibitor malonate regulates mitochondrial ATP-sensitive potassium channels: implications for ischemic preconditioning. Biochim Biophys Acta 2008;1777:882–889.

